# Turbulence in the Hippocampus: An Ansatz for the Energy Cascade Hypothesis

**DOI:** 10.1101/453506

**Authors:** Alex Sheremet, Yu Qin, Jack P. Kennedy, Yuchen Zhou, Andrew P. Maurer

## Abstract

Mesoscopic neural activity may play an important role in the cross-scale integration of brain activity and in the emergence of cognitive behavior. Mesoscale activity in the cortex can be defined as the organization of activity of large populations of neurons into collective actions, such as traveling waves in the hippocampus. A comprehensive description of collective activity is still lacking, in part because it cannot be built directly with methods and models developed for the microscale (individual neurons): the laws governing mesoscale dynamics are different from those governing a few neurons. To identify the characteristic features of mesoscopic dynamics, and to lay the foundations for a theoretical description of mesoscopic activity in the hippocampus, we conduct a comprehensive examination of observational data of hippocampal local field potential (LFP) recordings. We use the strong correlation between rat running-speed and the LFP power to parameterize the energy input into the hippocampus, and show that both the power, and the nonlinearity of mesoscopic scales of collective action (e.g., theta and gamma rhythms) increase as with energy input. Our results point to a few fundamental characteristics: collective-action dynamics are stochastic (the precise state of a single neuron is irrelevant), weakly nonlinear, and weakly dissipative. These are the principles of the theory of weak turbulence. Therefore, we propose weak turbulence as an ansatz for the development of a theoretical description of mesoscopic activity. The perspective of weak turbulence provides simple and meaningful explanations for the major features observed in the evolution of LFP spectra and bispectra with energy input, such as spectral slopes and their evolution, the increased nonlinear coupling observed between theta and gamma, as well as specific phase lags associated with their interaction. The weak turbulence ansatz is consistent with the theory of self organized criticality, which provides a simple explanation for the existence of the power-law background spectrum, and could provide a unifying approach to modeling the dynamics of mesoscopic activity.

## 1. INTRODUCTION

Hebb’s [1958] hypothesis that no psychological function can be attributed uniquely to any segment of cortex has the profound implication that cognition emerges from the coordination of activity across all spatial and temporal scales of the brain [Lashley, 1958, Allen and Collins, 2013]. The corollary is that understanding the brain begins with studying the the nature and role of different scales of brain activity. Temporal scales are typically more accessible to observations, defined based on the frequency structure of extracellular recordings (local-field potential, LFP), and assumed to be in a monotonic relation to spatial scales (lower frequencies correspond to larger populations [Buzsaki and Draguhn, 2004]).

The Fourier spectra of hippocampal LFP recordings cover a frequency range approximately from 0.05 Hz to 500 Hz, with spectra generally decaying as a power law. Two scales are readily identified. The high end of the spectrum, say,*ƒ* > 200 Hz^1^ represents the microscale, mainly occupied by action potential activity of single neurons, fast synaptic time constants, ion channel opening and closing, and heat dissipation (energy sinks)^2^. These processes are generated by microscopic neural units such as individual neurons, pairs of excitatory/inhibitory neurons, or small sequence. The processes occupying the low end of the frequency spectrum (say, *ƒ* < 60 Hz; theta rhythm and its harmonics e.g., Sheremet et al., 2016) are macroscopic in the sense that they encompass the several macroscopic segments of the brain. The theta rhythm is observed across several brain segments (hippocampus, entorhinal cortex, hypothalamus, prefrontal cortex, and others; e.g., Vertes and B., 1997, Siapas et al., 2005, Buzsaki, 2002), is associated with active exploration and REM sleep, and is assumed to provide the temporal structure for the organization of local networks [Green and Arduini, 1954, Green and Machne, 1955, Vanderwolf, 1969, Lisman and Idiart, 1995, Buzsaki, 2002].

In addition to these two scales, [e.g., Freeman, 2000b, Lubenov and Siapas, 2009, Patel et al., 2012, Muller et al., 2018, Zhang Honghui et al., 2018] cortical activity exhibits an intermediate, “meso” scale [e.g., Freeman, 2000b, Lubenov and Siapas, 2009, Patel et al., 2012, Muller et al., 2018, Zhang Honghui et al., 2018], which spans the frequency band 60 < *ƒ* < 200 Hz, a range of oscillations collectively referred to as the gamma rhythm [Buzsaki et al., 1983, Bragin et al., 1995, Belluscio et al., 2012, Lasztoczi and Klausberger, 2014, Schomburg et al., 2014]. Mesoscopic processes have been observed within brain regions associated with higher cognition, such as the neocortex and the hippocampus, that display a very specific laminar, highly isotropic and homogeneous (amorphous) structure^3^. The argument of the monotonic relation between temporal and spatial scales implies that the mesoscopic spatial scales are in the order of mm, which suggests that they represent *collective neuronal activity* (i.e., synchronized neuronal firing; Buzsaki and Draguhn, 2004)^4^. Many studies report several intriguing features of theta and gamma rhythms: during spatial exploration both theta and gamma respond to the intensity of activity as measured by rat speed with increased power (theta: Whishaw and Vanderwolf, 1973, Morris and Hagan, 1983; gamma: Chen et al., 2011, Ahmed and Mehta, 2012, Kemere et al., 2013, Zheng et al., 2015). Furthermore, it has been demonstrated that theta and gamma can couple as a function of task behavior [Bragin et al., 1995, Chorbak and Buzsaki, 1998, Chen et al., 2011, Ahmed and Mehta, 2012, Kemere et al., 2013, Zheng et al., 2015], and that coupling occurs in an energy dependent manner [Sheremet et al., 2018a]. Remarkably, the spatial structure of collective action spanning the theta-200 Hz frequency range appears to be that of propagating perturbations, i.e., waves [Lubenov and Siapas, 2009, Patel et al., 2012, 2013, Petsche and Stumpf, 1960, Muller et al., 2018]. These observations are particularly intriguing, because amorphous structures, with a dense and random network topology, are indeed the type of environment that would favor collective action over that of microscopic elements (signals coming from any specific connection would likely be “drowned out” by input arriving randomly through the random connections; furthermore, high synaptic failure rates between neurons are often quite high Freeman, 2000a, Buzsaki, 2006, English et al.,2017).

While mesoscopic collective activity could be explained away as a marginally-significant synchronization effect of strongly nonlinear microscopic units [Buzsaki, 2006], its strong relation with amorphous networks and intense behavior suggest that it might be in fact *the function* of this type of network. Freeman and Vitiello [2010] hypothesize, citing Lashley [1942], that mesoscale processes reflect the essential cognition step of abstraction and generalization of a particular stimulus to a category of equivalent inputs, “because they require the *formation of nonlocal, very large-scale statistical ensembles* (our emphasis)”^5^.

Taking this reasoning to its natural consequences, *we conjecture that cognition emerges as a function of the mesoscopic processes, and therefore has a fundamentally stochastic character*. This conjecture changes significantly the interpretation of observations. Rather than attributing special meanings to the activity of microscopic elements (like musical scores played by separate instruments), LFP recordings should be interpreted as local observations of a stochastic process representing collective action by mesoscale ensembles involving a large number of microscopic units: the precise state of a single unit is irrelevant.

Here, we examine some consequences of this change of perspective. Data collection and analysis procedures and detailed in Section 2. In section 3.1 we conduct a brief re-examination of basic elements of the stochastic analysis of rat hippocampal LFP observations, spectra and bispectra, in relation with rat speed. Despite some previous efforts [e.g., Wilson and Cowan, 1973, Wright and Liley, 1995, Freeman, 2000b, 2006, 2007, Freeman and Vitiello, 2010, Cowan et al., 2016]), a consistent theoretical approach to mesoscopic dynamics is missing. The purpose of the ansatz presented in section 3.2 is to provide a starting point for formulating such an approach^6^. The discussion in section 3.3 focuses on the weak turbulence description of collective action and its implications for the mesoscopic energy balance in the hippocampus. We demonstrate some capabilities of the theory by using the three-wave equations (a simplified, universal nonlinear interaction modeling framework) a to re-evaluate the significance of linear and nonlinear structure of hippocampal LFP recordings (section 3). The proposed theoretical framework is discussed in section 4. Some details of the algebra involved in the model derivation are given in section A.

## 2. MATERIALS AND METHODS

### 2.1. Subjects and Behavioral Training

Rat `r539♂-maurer` used in this study belongs to a cohort of 4-9 months old Fisher344-Brown Norway Rats (Taconic; see e.g, Zhou et al. 2018 for additional information about the cohort). The methods detailing the collection for these rats have been described in detail elsewhere [Zhou et al., 2018]. Briefly, following acclimation to the University of Florida colony, animals were trained to traverse a circular track for food reward (45mg, unflavored dustless precision pellets; BioServ, New Jersey; Product #F0021). During this time, their body weight was slowly reduced to 85% to their ad libitum baseline. Rats were surgically implanted with a custom single shank silicon probe from NeuroNexus (Ann Arbor, MI), designed such that thirty-two recording sites spanned across multiple hippocampal lamina.For probe preparation instructions and surgical methods, see [Vandecasteele et al., 2012, Zhou et al., 2018].

All behavioral procedures were performed in accordance with the National Institutes of Health guidelines for rodents and with protocols approved by the University of Florida Institutional Animal Care and Use Committee.

### 2.2. Neurophysiology

Following recovery from surgery, the rat was retrained to run unidirectionally on a circle track (outer diameter: 115 cm, inner diameter: 88 cm), receiving food reward at a single location. Following a few days of circle track running, the rats were trained to run on a digital-8 maze (121×101cm, L×W). During these sessions, the local-field potential was record on a Tucker-Davis Neurophysiology System (Alachua, FL) at ~24k Hz (PZ2 and RZ2, Tucker-Davis Technologies). The animals position was recorded at 30 frames/s with a spatial resolution less than 0.5 cm/pixel.

### 2.3. Spectral analysis3.1.2

The spectral analysis of the LFP in the current study was based on standard techniques used for stationary signals Priestley, 1981, Papoulis and Pillai, 2002. Descriptions of the stochastic estimators, meaning, normalization procedures, as well as how to interpret the bispectral maps, are given in Hasselmann et al. [1963], Rosenblatt and Van Ness [1965], Swami et al. [2001], Elgar [1987], Elgar and Guza [1985], Sheremet et al. [2016], and many others. Here, we remind the reader the main definitions and terminology.

Assume the LFP recordings *g(t)* and *h(t)* are realizations of zero-mean stochastic processes, stationary in the relevant statistics, with Fourier transforms *G*(*ƒ*_n_) and *H*(*ƒ_n_*), *n* = 1, …, *N*. The second and third order spectral statistics are estimated using cross-spectrum and bis-pectrum, defined as

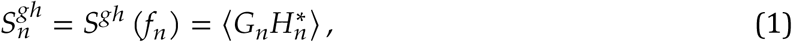

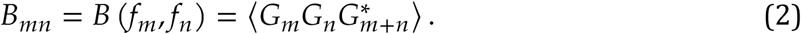

where the angular brackets denote the ensemble average, the asterisk denotes complex conjugation. We will generally omit the superscript when a single time series is involved. The diagonal 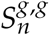 of the cross-spectrum matrix are power spectra. The coherence and phase lag of time series *g* and *h* are the normalized modulus and phase of the cross-spectrum,

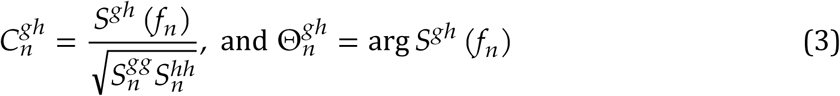

The cross-spectrum matrix provides information about the degree of correlation and phase lags for between different time series; spectra describe the frequency distribution of the variance of processes *g* and *h*, i.e., a complete characterization of the average linear structure of the Fourier representation.

The bispectrum provides information about the phase correlations between different frequency components of the same time series [e.g., Sheremet et al., 2016, Kovach et al., 2018]. The bispectrum is statistically zero if the Fourier coefficients are mutually independent, i.e., for a linear system, and will exhibit peaks at triads (*ƒ_n_,ƒ_m_,ƒ_n+m_*) that are phase correlated. The real and imaginary part of the bispectrum are related to the skewness 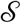 (e.g., positive skewness corresponds to sharp barrow peaks and flat troughs) and asymmetry 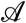 (e.g., positive asymmetry corresponds to the front of the wave being steeper than the back, similar to a saw-tooth wave) of the time series *g* through

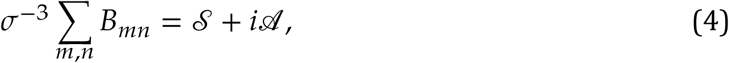

where *σ* is its standard deviation [Masuda and Kuo, 1981, Haubrich and MacKenzie, 1965]. The bicoherence *b_mn_* and biphase Φ*_mn_* are defined in way similar to the coherence and phase lag as the normalized modulus and the argument of the bispectrum, that is

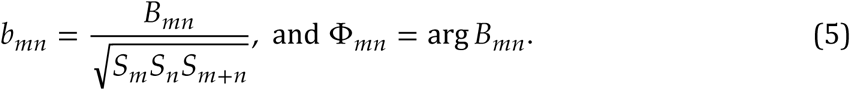

### 2.4. LFP power

Because the quantity measured by LFP observations is the potential of the electromagnetic field generated by the synaptic pulses, the spectral distribution of LFP variance (i.e., LFP spectral density integrated over some frequency interval) is proportional to the energy of the electromagnetic field per unit volume. Indeed, if the LFP time series *g*(*t*) is a zero-mean, weakly nonlinear stochastic process, stationary in the second order statistics [Priestley, 1981, Percival and Walden, 2009], one can show that the discrete Fourier representation has the property

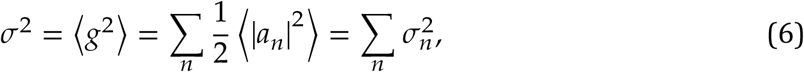

where 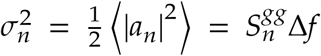 is the variance of Fourier mode *n, a_n_* is its amplitude, 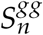 is the spectral density (equation 1), and Δ*ƒ* is the frequency band width. The mean energy per unit volume of an electromagnetic wave with the amplitude of the electric potential *g* is *w* = *εg*^2^,where ε is the dielectric constant of the medium [e.g., Pollack and Stump, 2002, Nolting, 2016]. The energy per unit volume of a stochastic electromagnetic field is therefore

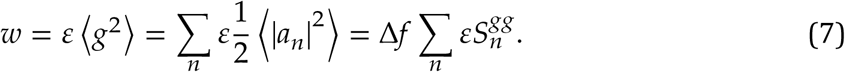

In other words, the amplitude squared of the Fourier components of the LFP are proportional to the energy stored in the unit volume by that particular Fourier component. A large number of algorithms have been developed for estimating for second- and higher-order statistics (power spectral density, bispectra, skewness, asymmetry, etc) of such processes. The analysis of the LFP in the current study was based on standard techniques used for variance-stationary signals [Priestley, 1981, Papoulis and Pillai, 2002] as previously described in Sheremet et al. [2016].

### 2.5. Numerical implementation

All data analysis was performed in Matlab® (MathWorks, Natick, MA, USA) using in-house developed code, as well as code imported from the HOSAtoolbox [Swami et al., 2001] for higher order spectral analysis.

Hippocampal layers were determined from the location of current sources and sinks derived on ripple and theta events, gamma power, and the polarity of the sharp-wave [Buzsaki, 1986, Buzsaki et al., 1986, Bragin et al., 1995, Ylinen et al., 1995, Lubenov and Siapas, 2009, Fernandez-Ruiz et al., 2017]. Details are given in Zhou et al. 2018.

The rat speed was calculated as the smoothed derivative of position. Raw LFP records sampled at 24 kHz (Tucker-Davis system) were pre-processed by applying a 2-kHz low-pass filter and divided into segments of 2048 time samples (approx. 1 second). Spectra and bispectra were classified by speed by averaging the speed over each of the 1-s LFP segments.

The dynamical and kinetic three-wave equations was integrated using the ODE solvers provided by Matlab®. The implementation is trivial, therefore, to save space, the codes are not provided. The authors will, however, gladly share them upon request.

## 3. RESULTS

### 3.1. Observations: stochastic measures of hippocampal activity

The monotonic relationship between rat speed and the power of collective action at theta and gamma scales [Morris and Hagan, 1983, Whishaw and Vanderwolf, 1973, Chen et al., 2011, Ahmed and Mehta, 2012, Kemere et al., 2013, Zheng et al., 2015] provides a parameterization of hippocampal activity in terms of animal running speed as well as energy input into the hippocampus (firing rates of hippocampal and entorhinal neurons increase with speed [McNaughton et al., 1983, Rivas et al., 1996, Shen et al., 1997, Hirase et al., 1999, Maurer et al., 2005, Kropff et al., 2015]). This phenomenon is easily observable for speeds roughly above 5 cm/s (e.g., figure 1). While low speeds, say < 3 cm/s may represent behavior uncorrelated to movement Zhou et al. [2018], higher speeds exhibit a strong correlation with hippocampal activity. Below we use this monotonic relationship both as a dependence of energy input into the hippocampus, and as a reference to the LFP power.

**FIGURE 1.**
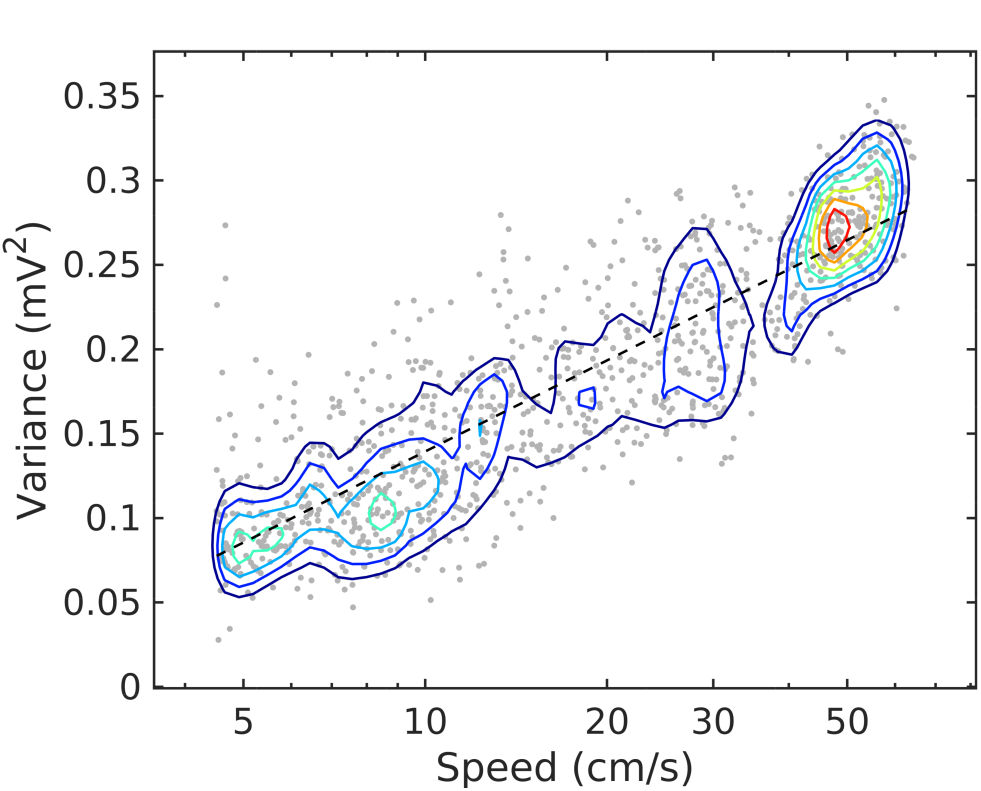
Joint probability density function for LFP variance (LM) and rat speed, for rat `r539♈-maurer`. The dashed line is a best-fit analytical expression *V* = *a*log_10_ *V* + *b*, with *a* = 0.18 and *b* = −0.04. Dots represent individual realizations (1-s time segments of the LFP recording, section 2.5). The relation between variance and speed follows the Weber-Fechner law of stimulus perception [Weber, 1860, Fechner, 1860].

For simplicity, we limit the data used here for illustration to a single representative source, rat `r539♂-maurer`. A discussion of the consistency of theses trends across the available, small (but growing) population of rats is presented elsewhere [Zhou et al., 2018, Sheremet et al., 2018].

#### 3.1.1. Variance spectra as a function of speed

Here, we examine estimates of spectral density of LFP traces recorded in the str. pyramidale (CA1.pyr), radiatum (CA1.rad), lacunosum moleculare (LM) and upper blade of the dentate gyrus (DG), indexed by rat speed.

Spectral densities show a weak, but significant variability as a function of speed and layer (figure 2). At lowest discernible speeds (*V* ≲ 3 cm/s), all hippocampal layers exhibit non-trivial baseline, lowest-variance spectrum, that can be characterized in general as having a power-law shape *ƒ^-α^* over the entire range of approximately 6 Hz to 300 Hz, with the exception of the DG layer, which exhibits a two-slope shape with a break point in the neighborhood of 50 Hz. In the CA1 layers the spectral slope is *α* ≈ 2 with slight variations (figure 2). In the DG, the slope of the lower frequency range *ƒ* ≲ 50 Hz visibly smaller, *α* ≈ 1. At high speeds (*v* > 15 cm/s), the theta rhythm and its harmonics dominate the lower frequency band of the spectrum, and low-power peak appears in the gamma range between 50 Hz and 120 Hz. Overall, the spectral slope of the spectrum decreases. The CA1 layers have similar slopes 1.4 < *α* < 1.7, with the DG layer again standing out at *α* ≈ 0.7. The power in the high-frequency tail of the spectrum (*ƒ* > 200 Hz) also increases significantly. In agreement with prior research [Bragin et al., 1995], the CA1.pyr layer shows the least energetic gamma range of all the layers examined.

**FIGURE 2.**
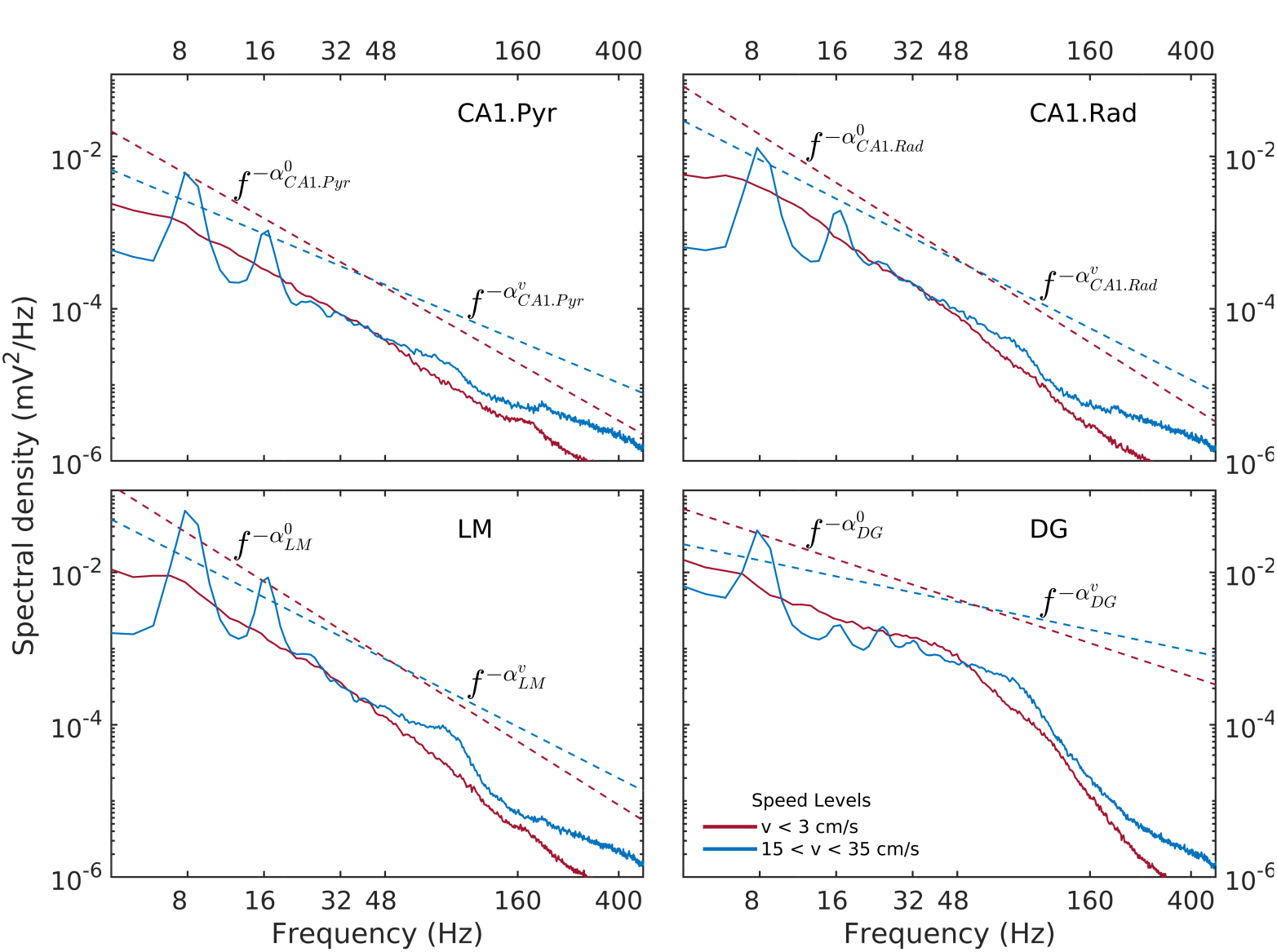
Power spectral density as a function of rat and hippocampal layer for low (red) and high rat speed (blue). Dashed lines represent power laws shapes *ƒ^-α^* that approximately match the observed spectra for 6 Hz < *ƒ* < 50 Hz. Low-speed spectral slopes are 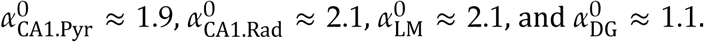 High-speed spectral slopes aree 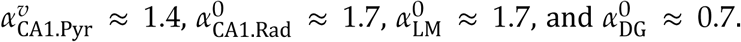 Data was produced from rat `r539♂-maurer`.

Details of the variability of hippocampal LFP spectra with speed are shown in figure 3 for seven speed intervals. According to figure 1, speed ordering is statistically equivalent to ordering by total LFP variance. The spectra are normalized by dividing them by *ƒ^-α^*, using the high-speed slopes 1.4, 1.7, 1.7, and 0.7 for CA1. Pyr, CA1.Rad, LM, and DG respectively. The normalization reduces the spectra to ≈ 1 in the frequency range where it agrees to the power law, and highlights spectral peaks. Although the ordering parameter is total variance, the spectra show a remarkable monotonic ordering in all frequency bands, with the exception of the highest two speed intervals, where the evolution stagnates and perhaps reverses slightly. The normalization re-scaling highlights several features of the evolution as a function of speed. The spectra in all layers tilt as energy increases (e.g., the LM slope changes from 2.1 to 1.7). Theta and its harmonics become dominant in the low-frequency range: four peaks at multiples of 8 Hz may be seen clearly in the CA1.Pyr and DG spectrum, perhaps three in the CA1.Rad and LM spectra. Also remarkable is the peculiar way the gamma range evolves: rather than developing some broad peak in 50-120 Hz range, the gamma range growth seems to result from the *s* ≈ 1 domain (where *s* is the normalized spectrum) progressively extending into the higher frequency range, while a “bump” develops in the neighborhood of *ƒ* ≈ 100 Hz. This evolution is suggestive of a “front” of energy that propagates “against some resistance” toward higher frequency. This type of evolution may be seen in all layers. Although one might expect slight differences between acceleration and deceleration states for the same speed, these are negligible (data not shown). Therefore, the classification by speed does not differentiate between acceleration sign: the transition, say, between 10 cm/s and 20 cm/s may includes both acceleration and deceleration. Thus, the transition process described by figures 1, 3 (and also below) is reversible, therefore quasi-stationary: all states described by these spectral may be assumed quasi-equilibrium states (in other words, imagining this evolution as a number of time steps, the transition between different steps is slow enough to allow the system to reach equilibrium and every step). This is consistent with the ability of the rat to control its speed. We will return to these ideas below.

**FIGURE 3.**
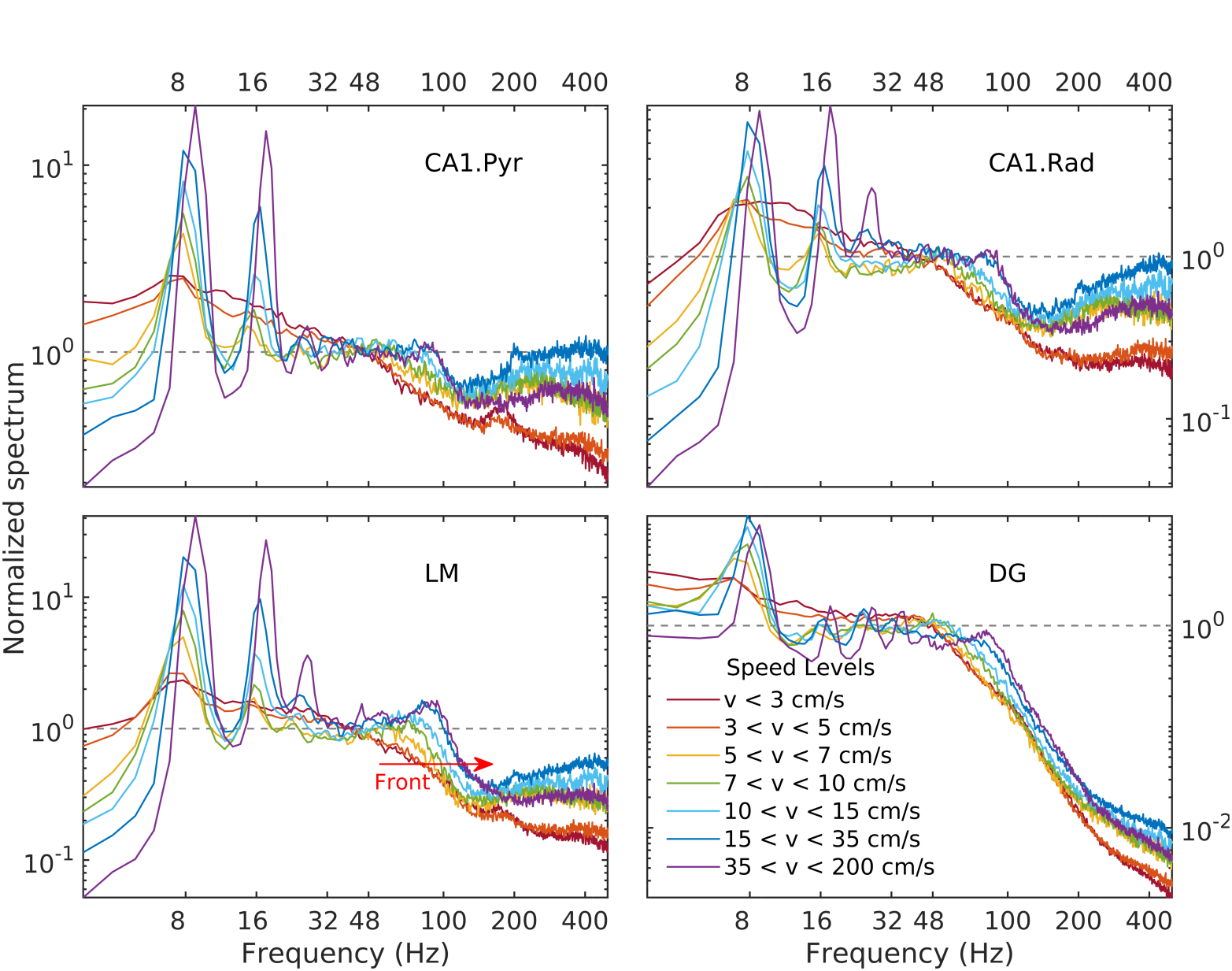
Normalized LFP spectrum estimated for CA1.pyr, CA1.rad, LM, and DG layers as a function of speed. Frequency spectra are normalized by dividing them by the corresponding power law, i.e., 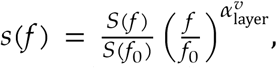 where *S* is the power spectrum, *s* is the normalized spectrum, and *ƒ*_0_ = 48 Hz. The evolution of the LM gamma band has the appearance of a spectral “front” moving to the right (red arrow), as it steepens and develops a peak. A similar behavior may be seen in all layers.

#### 3.1.2. Higher-order spectra as a function of speed

Higher-order spectra provide information about cross-frequency coupling. The bispectrum, the lowest order (and hence, the most “accessible” such estimator) has been used for a long time in wave dynamics Hasselmann et al., 1963, Rosenblatt and Van Ness, 1965, Coppi et al., 1969). The relationship between its structure and third order statistics of the time series is well understood (e.g., Haubrich and MacKenzie, Masuda and Kuo, Elgar; for bispectral definitions and terminology, see section 3.1.2). Although bispectral analysis is not common in neuroscience, recent work [Kovach et al., 2018] has shown that similar, widely used estimators for phase-amplitude and amplitude-amplitude coupling are, in fact, particular implementations of bispectral estimators (containing additional restrictive assumptions that make them susceptible to misinterpretation; e.g., Hyafil, 2015). To save space, we discuss only low (*v* < 10 cm/s) and high (*v* > 35 cm/s) speed levels; intermediate levels (not shown) represent a relatively smooth transition between these two limits. Because of the substantial difference in dynamics between CA1 and DG, we confine our discussion of bispectra to the LM layer.

As previously reported [Sheremet et al., 2016], bispectral estimates also show significant variability with rat speed. Low-speed bispectra are statistically zero (Gaussian) overall (figure 4). The bicoherence exhibits a weak peak at (*ƒ_θ_, ƒ_θ_*, 2*ƒ_θ_*), corresponding to the phase coupling between theta (*ƒ_θ_* ≈ 8 Hz) and a relatively broad band around its 2*ƒ_θ_* harmonic (see arrow, figure 4a, bicoherence). The phase of the harmonic band is in part in opposition and part in quadrature with theta (see transition red to blue in figure 4, biphase), thus contributing overall to negative skewness of the LFP (peak is negative in 4a, skewness), but positive asymmetry (4a, asymmetry; see definitions in section).

**FIGURE 4.**
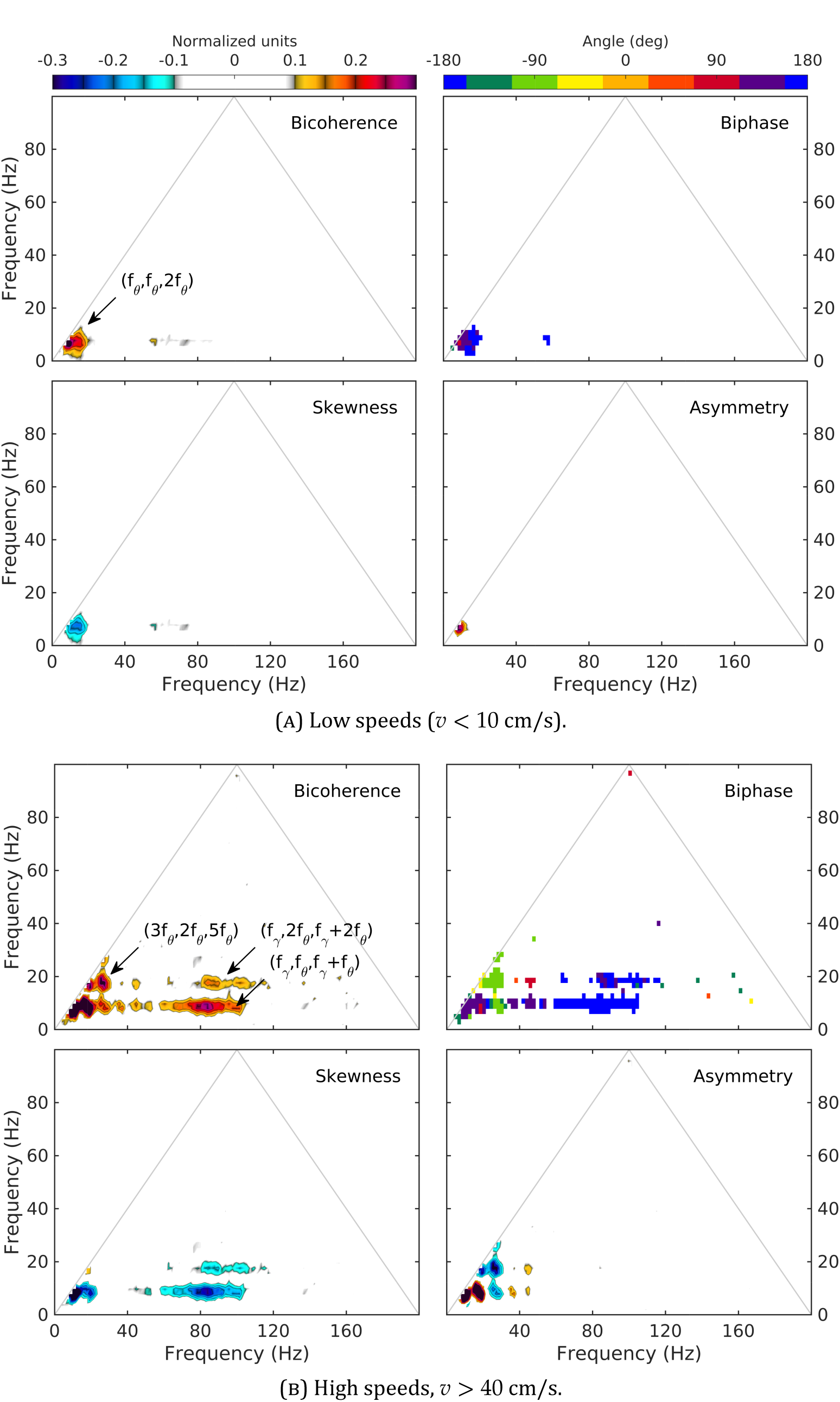
Normalized bispectrum (equation 5) for the LM layer. The bicoherence is blanked below 0.1 (with 300 DOF, zero-mean bicoherence is <0.1 at 95% confidence level; Haubrich and MacKenzie, 1965, Elgar and Guza, 1985)

In contrast, high speed activity show rich phase-coupling structures, involving theta and gamma (figure 4b). One may separate two frequency regions corresponding to the coupling between theta and its harmonics, and theta and gamma. The picture of the coupling between theta and its harmonics agrees with the spectral evolution (figure 2) and previously reported results

[Sheremet et al., 2016]. The bicoherence exhibits significant levels of phase coupling, reaching as high as (5*ƒ*_*θ*_;,*ƒ*_*θ*_,6*ƒ*_*θ*_), (3*ƒ*_*θ*_,2*ƒ*_*θ*_,5*ƒ*_*θ*_),and (3*ƒ*_*θ*_,3*ƒ*_*θ*_,6*ƒ*_*θ*_). The relationship between harmonics and theta is quite diverse, with some harmonics contributing to LFP skewness, and others only to LFP asymmetry. For example (figure 4b, skewness and asymmetry), the coupling (*ƒ*_*θ*_, *ƒ*_*θ*_, 2*ƒ*_*θ*_) generates both negative skewness and positive asymmetry, while (2*ƒ*_*θ*_,2*ƒ*_*θ*_,4*ƒ*_*θ*_) results in negative asymmetry only. Theta-gamma coupling is prominent at high speed, engaging theta harmonics, and contributing strongly to negative skewness (figure 4b).

#### 3.1.3. Summary of observations

Our observations show that sorting LFP epochs by speed provides an efficient classification device that produces remarkably well ordered spectra and bispectra. As summarized in figure 5, with increased speed the total LFP power increases, as well as the power in the theta and gamma bands. The power in the theta band grows overall by a factor of 4 at a relatively steady rate, while gamma grows by about a factor of 2 and seems to plateau. Phase coupling (as measured by the bicoherence integrated over the theta/harmonics and theta/gamma frequency domains) shows a steady growth, accelerating at high speeds. Spectral slopes (*α*) decrease, consistent with an accumulation of energy in the gamma range; and phase coupling involving theta and gamma increases.

**FIGURE 5.**
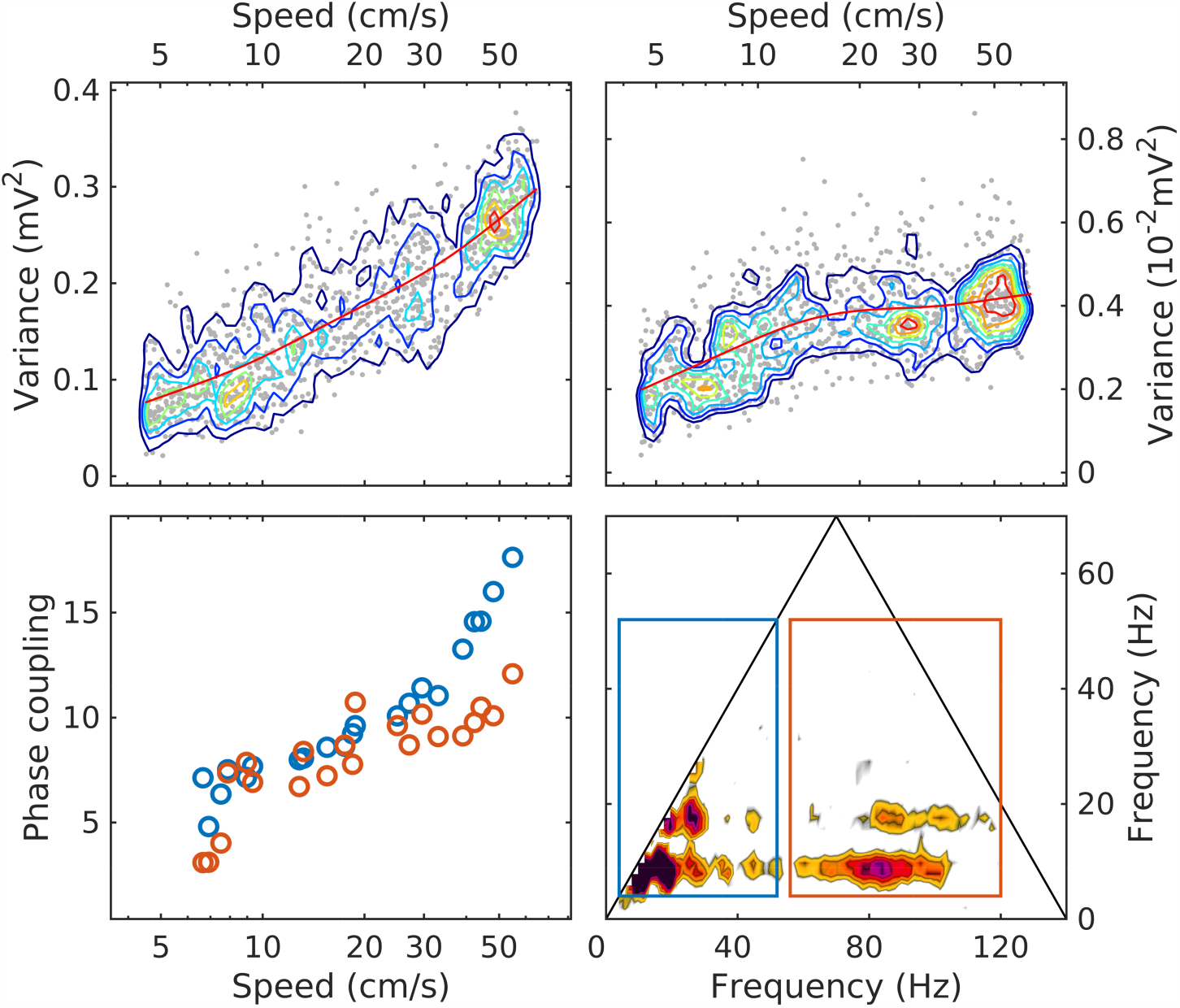
The evolution with speed of LM theta power (upper left); gamma power (upper right; note the different units); phase coupling of theta and its harmonics (lower left, blue); phase coupling of theta and gamma (lower right, red). The phase coupling measure is the bicoherence integrated over the are of the rectangles shown in the lower-right panel (same colors as in lower-left panel). Because the bicoherence is normalized, the units of the phase coupling measure are arbitrary. In upper panels, the red line is a moving average, included to highlight the evolution trend.

The process of evolution with speed is reversible. In “reverse” the process appears to converge as *v* → 0 toward a limiting, “background” state. Ignoring for now other forms of activity not taken into account here, the background state represents “inactive” behavior, whereas the active state would be associated with significant speed levels. This hypothetical background has some remarkable properties. Although its spectrum decays faster than any active-state spectrum (higher slope *α*), it still contains significant power (in figure 1, about 1/4 of the most active state). In addition, the LFP is nearly Gaussian, i.e. exhibits overall no phase coupling (ignoring the weak theta signal).

## 3.2. Ansatz: mesoscopic wave turbulence

The goal of this study is to provide an interpretation of the observations. Our stochastic conjecture collective implies that mesoscopic collective action is the meaningful signal in the hippocampus, and one should interpret the spectral and bispectral trends seen in observations as the expression of collective action dynamics.

### 3.2.1. Why an ansatz?

Mesoscopic collective action has been observed [e.g., Freeman, 1975, Lubenov and Siapas, 2009], manifesting in some cases as propagating waves [Lubenov and Siapas, 2009, Patel et al., 2012, 2013, Zhang Honghui et al., 2018, Muller et al., 2018]. Collective action is demonstrably weakly nonlinear (section 3.1.2), and also intrinsically stochastic, because it is macroscopic with respect to microscopic neural units (neurons, or neural micro-circuits), hence the precise state of a single neural unit must be irrelevant for the mesoscopic process. Observations of waves propagation suggest that, at least at some scales, mesoscopic collective action experiences negligible dissipation. That is, as opposed to microscopic dissipation (transfer of synaptic energy into other forms), the theta and gamma patterns do not show strong dissipation (that is, waves can propagate, retaining identity within the hippocampus).

Not much is known, however, about the spatial-temporal structure of collective action; about the mechanisms govern its dynamics; about its role in the energy balance of the global brain, or how it supports cognition (if the stochastic conjecture is meaningful). Modeling efforts [e.g., Cowan et al., 2016, Wilson and Cowan, 1972, 1973, Wright and Liley 1995] have been sparse and seem to lack a comprehensive, larger-physics context. It is important to note that, because of the scale separation, mesoscopic and microscopic dynamics are different, therefore the wealth of knowledge accumulated about microscopic physics cannot be directly extended to mesoscopic processes. While the whole inherits some of the properties of the parts, it will have its own physical laws and dynamics. The notion that theory is scale-dependent is common in physics: macroscopic systems are characterized by state variables and laws that are typically inaccessible directly to microscale theories. For example, the characteristics of molecules does not describe the difference between ice and liquid water. A relation exists between the micro-and macroscales, but it is subtle, mediated by averaging operators, and often result in surprising consequences. An example is Boltzmanns celebrated *H*-theorem, which introduces the entropy (*H*) as a new state variable, and elucidates the process through which time-reversible microscopic dynamics begets the macroscopic irreversibility of evolution toward equilibrium [Gibbs, 1902, Khinchin, 1949, Toda et al., 1983, Pathria and Beale, 2011]^7^.

Despite some previous efforts [e.g., Wilson and Cowan, 1973, Wright and Liley 1995, Freeman, 2000b, 2006, 2007, Freeman and Vitiello, 2010, Cowan et al., 2016]), a consistent theory of mesoscopic dynamics is missing. The purpose of an ansatz is to provide a starting point for formulating such a theory. In the least, it will suggest a new perspective for the interpretation of observational data, and may generate verifiable inferences about mesoscopic physics.

### 3.2.2. Turbulence

Consider an energy-conserving, physical system that is large enough to cover a range of scales (figure 6, left). If the system is nonlinear, scales interact and energy may be exchanged across scales. Assume that energy is fed into the system at large scales, and dissipation processes are most active at small scales. If the energy source and sink are well separated in scale, there exists an intermediate- scale (mesoscale) domain, called inertial range, that is largely free of sources and dissipation. In the intermediate scale, the only energy process is an energy flow (flux) from source in the direction of the sink.

**FIGURE 6.**
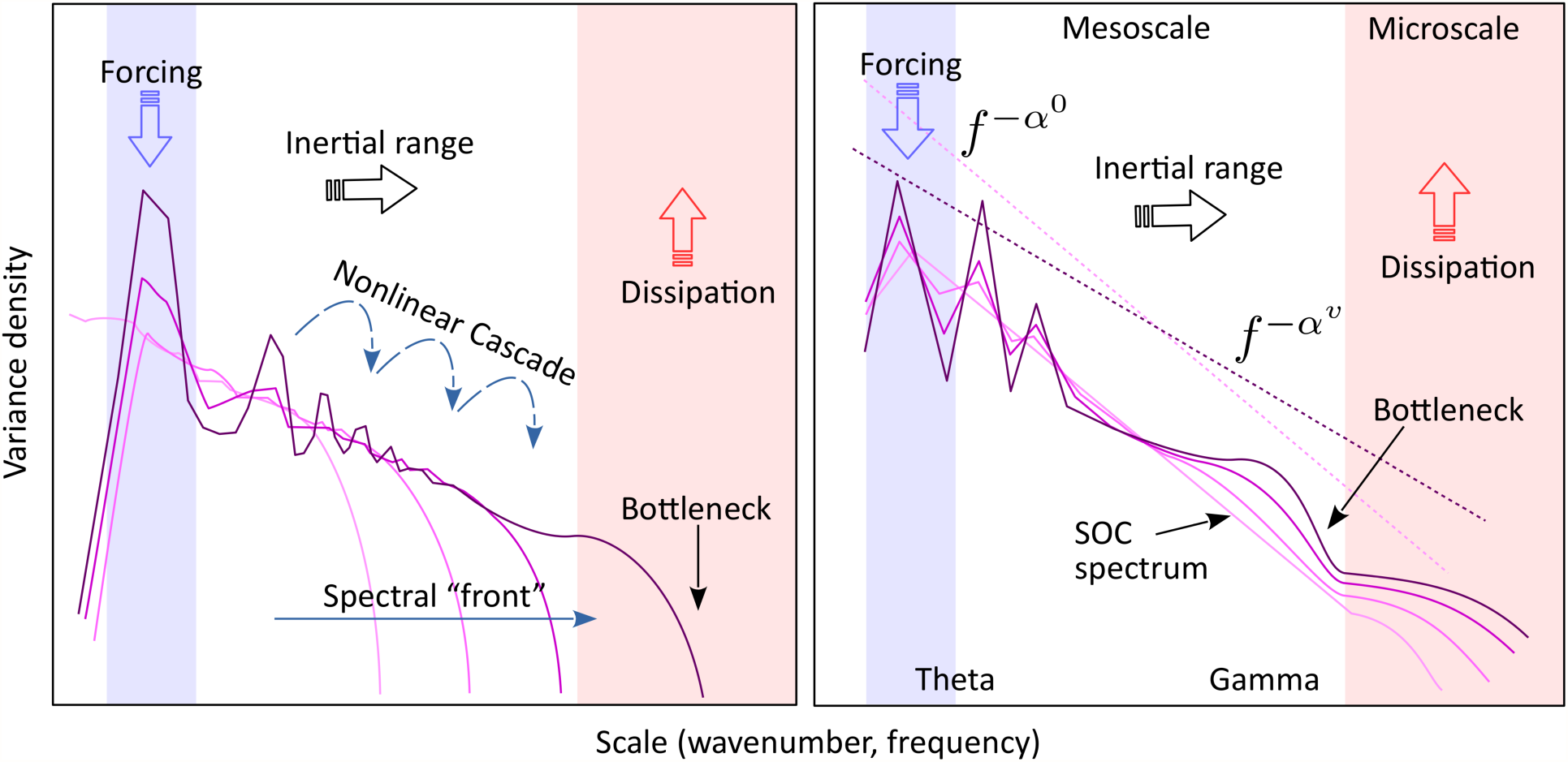
Left: The turbulence model. Energy is introduced into the system at the forcing scale (blue) is separated from the dissipation scale (red) by a “transparent” inertial range (white), largely free of forcing and dissipation. Nonlinear interactions generate a cross-scale energy flux (cascade) from the source to the sink. Gray curves show the possible evolution of the spectrum toward stationary state, if initially the inertial range contains no energy. The stationary spectrum (purple) corresponds to a constant spectral flux of energy across the inertial range. A “bottleneck” in the spectral flux capacity at small scales may cause an accumulation of energy (spectral bump) at larger nearby scales. The axes are in logarithmic scale. Right: A schematic of the observed spectral evolution, interpreted in a way similar to turbulence. The light-colored spectrum is the background, corresponding to low activity (speed), and representing the self-organized critical (SOC) state. Increasingly dark lines represent the spectral shape at increasing intensity of activity (speed). The low-frequency peaks represent the theta rhythm and its harmonics (marked collectively as “theta”). A feature similar to the turbulent spectral front is observed in the gamma range. A spectral bump similar to a bottleneck (e.g., Lvov et al., 2007, Meyers and Meneveau, 2008, Proment et al., 2009, Nazarenko, 2011, see note in the text) is observed in the high-frequency gamma range.

The energy balance of the system is controlled by the strength of source and sink characteristics, and the flow (flux) through the inertial range. Assume that the input rate at the source is constant, and the system has initially zero energy. The energy at the source scales will increase in time. As the energy of a given scale increases, the effectiveness of nonlinear exchanges across scales increases, and the energy introduced into the system flows downscale. Eventually, the energy flow (nonlinear cascade) reaches the dissipation (sink) scales, where it is removed from the system. The stationary state will occur when the dissipation rate matches the input rate, and will be characterized by a constant flux of energy across the inertial range^8^. The cross-scale energy flux is Richardsons [1922] turbulent cascade. In hydrodynamic turbulence the stationary spectrum follows a power law, the celebrated Kolmogorovs [1941] “five-thirds” law *k*^−5/3^, where scales are represented by the wavenumber *k*.

### 3.2.3. Mesoscopic turbulence on an active network

This turbulence model could be useful for collective action dynamics, *provided that*the mesoscale may be identified with the inertial range; or, equivalently if nonlinearity dominates dissipation at mesoscale. However, while observations seem to support this assumption (see discussion in section 3.1.3), it contradicts the well-known strongly-dissipative character of microscopic neuronal dynamics^9^. A further complication is that the turbulence model as described in figure 6 differs from observations (e.g., figure 2) at least in one important way: the live hippocampus never exhibits a zero energy state. Clearly, these points deserve some discussion.

We propose that the amorphous hippocampus structure constitutes an “active” network, that functions by chain reaction. From the stochastic mesoscopic perspective, neuron activity is meaningful only on average, and as a contribution to the overall network energy balance. Therefore, the “mesoscale neuron” is a microscopic device that is continually supplied with energy, which it releases in explosive bursts, activated by a threshold type of trigger. Bursts are followed by a “recharge” (refractory) period during which the energy reserve is replenished. Most of the burst energy is lost (e.g., as generating the electromagnetic field - LFP recordings measure in fact lost energy!), but a fraction of it is channeled through network connections to other neurons, in service of triggering additional bursts elsewhere. Therefore, while microscopic energy sources and dissipation are essential ingredients for an active system, *the “meaningful” activity of the system is driven by the energy that is channeled through the network* connections. In other words, collective action may be thought of as “waves” of channeled energy that achieve triggering values in the areas where they propagate. We will refer to this as “trigger” energy. Because trigger energy is replenished by each burst, it seems reasonable to assume it is subject to only weak dissipation^10^. Note that the definition of collective action as “trigger” energy also implies that its source is likely different from the source of energy of the microscopic network unit (neurons and such).

The active-network concept is consistent with non-zero background activity (background spectrum, section 3.1). Trying to propagate a wave of collective action through an active system at complete rest, i.e., when the “trigger energy” being channeled though the system is zero, is not “optimal” in the following sense: a relatively large amount of “trigger energy” is required; a large number of neurons will be triggered simultaneously, therefore driven simultaneously into refractory state, rendering large portions of the system temporarily inoperable; and the propagation of such collective action might not be sustainable, as bursts might not provide enough trigger energy. However, if a background level of trigger energy is available to maintain a large majority of neurons just below the burst threshold value, generating collective action would require “minimal” amounts of excess energy. The best way to establish such background level might be by random excitation of individual neurons; a large number of neurons (mesoscopic character of the collective action) would insure that there are enough neurons available to propagate the signal. One may note here a strong affinity with ideas such as self-organized criticality [e.g., Bak et al., 1988, Beggs and Plenz, 2003, Beggs and Timma, 2012, Pruessner, 2012] and metastability [Tognoli and Kelso, 2014]. The featureless, power-law, Gaussian background spectrum we observe (figure 2) is consistent with the “edge of chaos”, self-organized critical background state for optimal transmission of collective action.

We postulate that mesoscopic collective processes are weakly-nonlinear and weakly-dissipative trigger perturbations of a critical-type equilibrium state of the active neural network. The source of energy for collective action is different from that which powers microscopic units, may well be provided at the large scale end of the spectrum. Thus, collective action dynamics seem to satisfy basic conditions for the turbulence ansatz.

Figure 6, right panel, shows a schematic of the observed evolution of the LFP spectrum with speed. After accounting for the presence of a background spectrum, the resemblance with the turbulence model (6, left) is striking. The similarity suggests that theta is the source of energy for collective action, and microscopic processes are as the main energy sink, while the mesoscale acts largely as an inertial window that allows for a cross-scale energy flow from source to sink. The development of the gamma peak, similar to the bottleneck effect in hydrodynamics, may be interpreted as the existence of a transitional scale right above microscopic, that has a limited energy-flux ability, thus causing an accumulation of energy at intermediate scales. The observation that spectral evolution with speed is reversible (in other words, that spectra at each speed represent an equilibrium state) is also remarkable, because it is consistent with Kolmogorov stationary spectra of turbulence.

## 3.3. The weak turbulence model

The turbulence ansatz may now be formalized. Because collective action is approximated as conservative, we shall use as a starting point the assumption that collective action dynamics is Hamiltonian. We stress that, as a starting point, the hypothesis that nonlinearity dominates dissipation at mesoscale is essential and sufficient. The Hamiltonian assumption is not needed; we use if for convenience, because it allows for introducing weak turbulence principles in a general form, without having to reference the specifics of the system.

Here we give only the a brief elementary discussion of the basic equations that govern weak turbulence. The principles of the weak turbulence model, some derivation algebra, and some results are further given in appendix A. We caution, however, that this description is oversimplified, retold version, and that the full theory is much richer and complex. We encourage the interested reader to consult the original sources, written by the fathers of the theory: the comprehensive monograph Zakharov et al. [1992], excellent review papers by Zakharov, 1999, Newell et al., 2001, Newell and Rumpf, 2010, and the account by Nazarenko [2011], that includes more recent results.

### 3.3.1. Dynamical equation

The evolution equation for the Hamiltonian system has the universal form (see appendix A, also the remarks below)

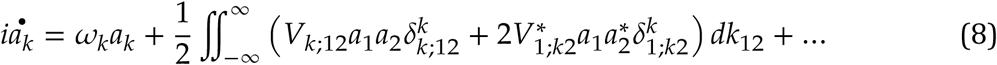

Equation 8 is written for the Fourier transform *a*(*k, t*) of a canonical variable that defines the mesoscale system (for details, see appendix A.2). A Fourier mode identified by its wavenum-ber *k* and frequency *ƒ*(*k*), (we assume that the dispersion relation 24 has a single root). The rest of notations are: *ω*(*k*) = 2*πƒ*(*k*) is the radian frequency; 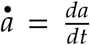; *V*_*k*;12_ = *V*(*k*;*k*_1_,*k*_2_) is the interaction coefficient; and *δ* is the Dirac delta function. We use the shorthand notation *σ*_1,2_ = *σ* (*k*_1,2_), *σ*_*k*_ = *σ*(*k*), and 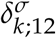 where *σ* is some quantity depending on *k*; and also *dk*_12_ = *dk*_1_*dk*_2_. For simplicity we limit the discussion quadratic nonlinearity (right-hand side terms of the form *a*^2^) terms, but a full description might require including cubic (*a*^3^), and possibly higher-order nonlinearity (e.g., the nonlinear Schrodinger equation, Newell, 1985; or the Ginzburg-Landau equation Ermentrout, 1981, Cross and Hohenberg, 1993, Passot and Newell, 1994, Ermentrout et al., 1997)

Two equations for the modulus *b* ≥ 0 and phase *θ* of *a*(*k,t*) are obtain substituting *a*(*k,t*) = *b*(*k,t*)*e*_*iθ(k,t)*_ into equation 8

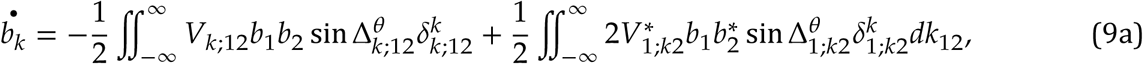

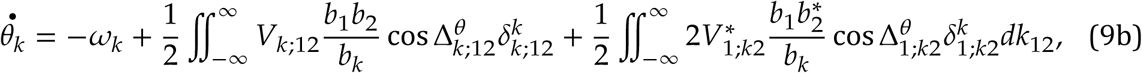

where we used the notation 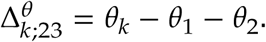

*Remarks*. As in thermodynamics, mesoscopic state variables are different from microscopic ones. For the hippocampus, such variables may be the average ratio of neurons activated per unit area and unit time [e.g., Cowan et al., 2016, Wilson and Cowan, 1972,1973, Wright and Liley 1995, Robinson et al., 1997]. Mesoscopic variables are related to microscopic ones in the same way as, for example, macroscopic pressure is related to microscopic forces resulting from collisions of individual molecules with a surface.

Temporal (frequencies) and spatial scales (wavenumbers) are related through the dispersion relation 24, therefore only one of the parameters *k* and *ƒ* is independent. The choice of the independent parameter is arbitrary, because the dispersion relation is invertible. The *ƒ*(*k*) representation resolves the spatial structures and yields a time-evolution equation. The *k*(*ƒ*) representation resolves the time structure and is appropriate for time-series analysis (e.g., LFP recordings). Therefore, equations are written here in *ƒ*(*k*), but observational data is discussed in the *k*(*ƒ*) representation. Below, the concepts of “frequency”, “wavenumber” “mode” and “scale” will be treated as equivalent and interchangeable.

Equation 8 is called dynamical equation. If the Hamiltonian 30 is identified with energy, the quantity |*a*|^2^ has the physical dimensions of action (energy×time). The form of equation 8 in universal in the sense that the details of the physics of the system are contained the coefficients *ω*_*k*_ and *V*_*k*;12_ only.

Equation 8 describes the nonlinear interaction of mode *k* with the pair of modes (*k*_1_,*k*_2_). A triplet of interacting modes (*k,k*_1_,*k*_2_) is called a “triad”. The strength of the interaction depends on the interaction coefficient *V*_*k*;12_. The factor 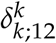, resulting from the orthogonality of the Fourier representation (equation 23), is a selection criterion: interacting modes satisfy the equation

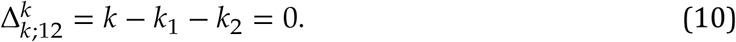

It is useful to think of equation 8 in a discretized form, e.g., replacing the integrals by sums. A schematic representation is shown in figure 7. Equations 9 show that nonlinear interaction result in both amplitude and phase evolution. The effectiveness of nonlinearity depends on modal amplitudes, on the interaction coefficient, and (importantly) on the phase mismatch 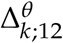 (see also section 3.3.2 below). If 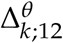 is large, the nonlinear term oscillates fast and the effect small; if 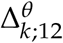 is small, the nonlinear term preserves sign over longer periods of time and the effect is significant. If 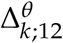, the contribution is maximal.

**FIGURE 7.**
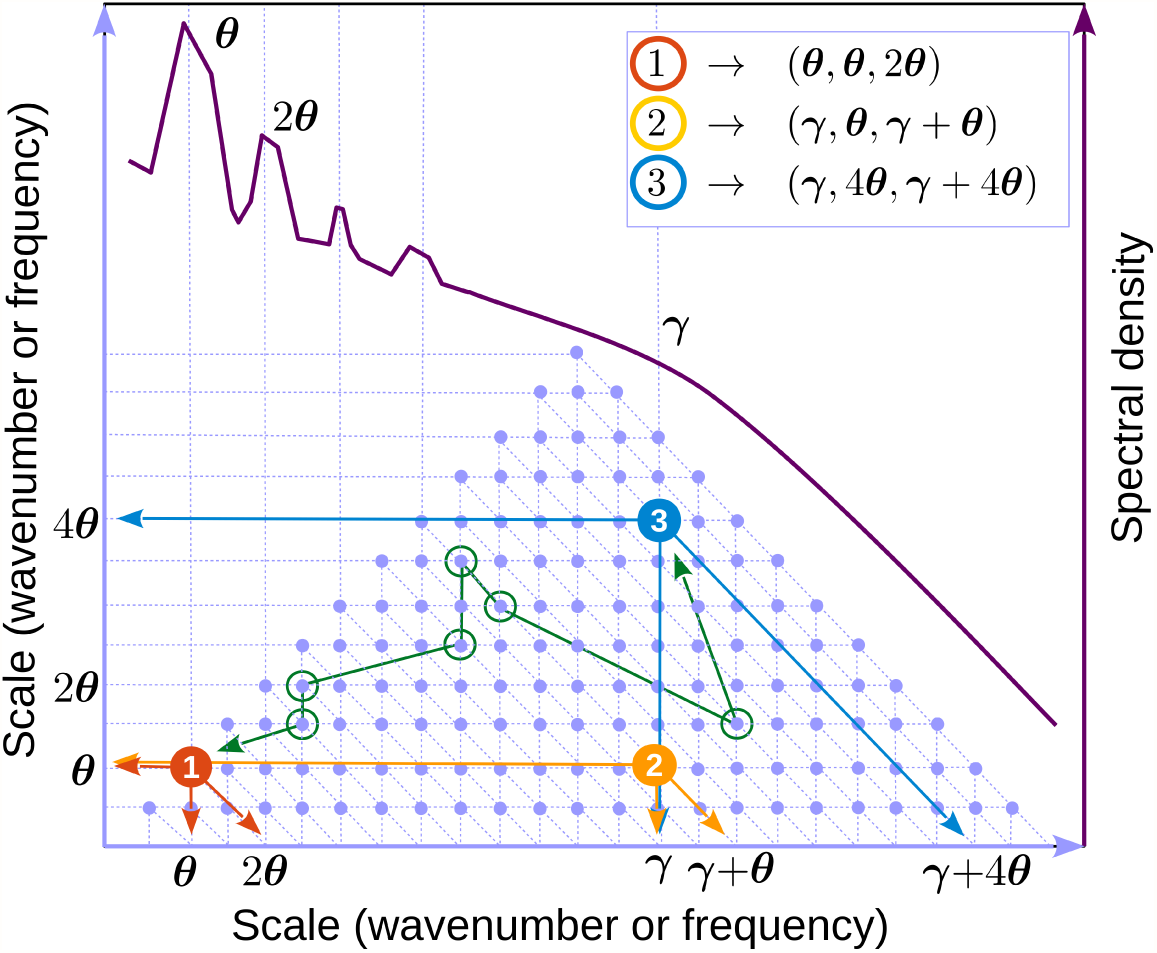
Discrete representation of interacting triads (equation 8). Because *k* and *ƒ* are interchangeable, we refer to either as “scale”. Light blue dots represent triads. The The axes represent scale 1 and 2 in the triad (e.g., *k*_1_ and *k*_2_ in equation 10). The third interacting mode (*k* in equation 10) may be found by as the intersection of the second diagonal passing through the triad with the horizontal axis (e.g., arrows in triad 1, red dot). One can easily check that the triangle of blue dots represents all possible triads for the scale interval shown (hence the triangular shape of the bispectrum). A schematic of a LFP spectral shape (dark red) is used to indicate possible locations for the theta (*θ*) and gamma (*γ*). Example of triads involving theta and gamma are identified by colored dots (compare with the annotations in figure 4b). Green circles mark an example (arbitrarily-chosen) of a chain of triads connecting triads 1 and 3. The order of connection is marked with a green line. One can check that each pair of consecutive circles share one mode. All triads in the spectrum are connected by many such chains.

The nonlinear contributions to modal phase evolution (equation 9b) have same properties as those in the amplitude equation 9a, with the important difference of *b*_*k*_ appearing at the denominator. If *b*_*k*_ → 0 (mode *k* has small amplitude) the nonlinear phase component becomes arbitrarily large and dominates the total phase. Therefore, the phase of low-amplitude waves is “dictated” by nonlinear forcing.

The dynamical equation 8 is deterministic: maybe integrated exactly to obtain the state of the system at time *t* if the initial value of amplitudes *a*(*k,t*_0_) are known (*t* > *t*_0_). Initial phases are difficult to measure in practical applications, but a description of the “average” behavior of the system, independent of the initial phases, can be readily constructed by integrating equation 8 many times with different sets of initial phases and averaging the results.

### 3.3.2. The kinetic equation

The last remark above suggests averaging the equation itself, rather than averaging the solutions. The averaging procedure and the associated closure problem are summarized in appendix A.3. Briefly, averaging introduces a hierarchy of averages of amplitude products (correlators), such as 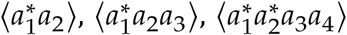, and so on, where the angular brackets denote the ensemble average. Assuming spatial homogeneity implies that

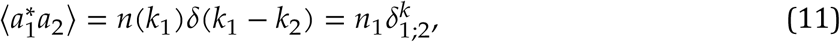

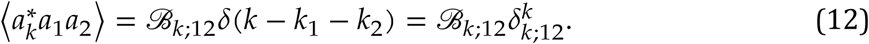

The quantity *n*(*k*) represents the action density, or “occupancy number”, or “number of particles” (by analogy with quantum mechanics). In statistics *n* and *ℬ* are called “spectrum” and “bispectrum”, respectively. The quasi-Gaussian assumption (equation 35) results in a system of two coupled equations for the spectrum and bispectrum (section A.3)

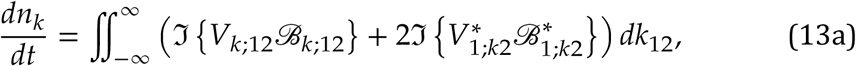

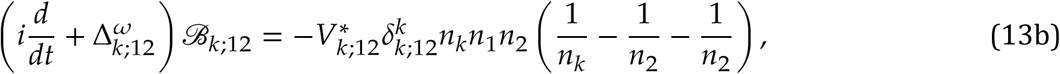

. Assuming additional long-time regularity conditions (section A.5) reduces the system to single equation called the *kinetic* equation

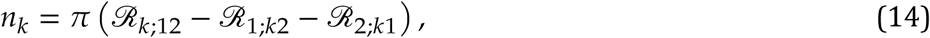

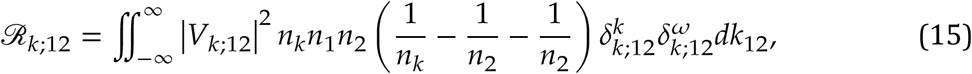

where

*Remarks*. Equation 13a highlights the dynamical significance of the bispectrum as the nonlinear forcing in the evolution of the spectrum. If the bispectrum cancels, nonlinear interactions cancel and the system is effectively, *on average*, linear.

The kinetic equation 14 or the more general system of equations 13, represent a stochastic, ensemble-averaged description of the system 8. Kinetic equations of the type of equation 14 were introduced in statistical mechanics by Boltzmann [e.g., Boltzmann, 1872, 2003, Alexeev, 2004], and are powerful tools in the study of multiple-scale system.

Equation 14 states that in the long-time limit the only interactions that are effective are due to triads that satisfy the *resonance* conditions imposed by the factors 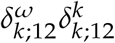, i.e., satisfying the conditions,

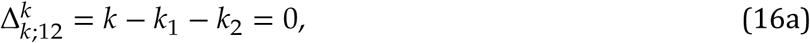

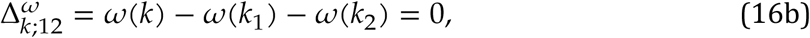

equivalent to the “maximal” effectiveness of nonlinear interaction (see discussion of equations 9). Whether or not equation 8 has resonant triads depends on the linear properties of the physical system. The resonance conditions play an important role in the stochastic theory [e.g., Zakharov et al., 1992, Nazarenko, 2011, Anenkov and Shrira, 2018].

### 3.3.3. Stationary spectra

Stationary spectra 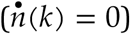 are of importance for systems whose evolution is a quasi-equilibrium process. Equation 14 has two classes of stationary solutions (e.g., Nazarenko, 2011, see also section A.7).

The Rayleigh-Jeans (RJ) class of spectra comprises the stationary solutions *k*^−1^, *ω*^−1^, and (*C*_*ω*_*ω* + *c*_*k*_*k*)^−1^, that obviously cancel the integrand in equation 14, and correspond to zero exchange across scales (spectral fluxes *ℱ*_*q*_ (*k*) ≡ 0, equation 43d). Therefore RJ spectra correspond to thermodynamic equilibrium and equipartition of momentum (*nk*) and energy (n*ω*). Realistic, non-isolated systems do not typically reach thermodynamic equilibrium, therefore RJ spectra are not important for the weak turbulence ansatz, which includes sources and sinks as essential elements.

The Kolmogorov-Zakharov (KZ) spectra are different class of stationary spectra, that correspond to a constant spectral flux *∂*_*k*_ *ℱ*_*q*_ (*k*) across the inertial range. They were derived for equation 14 by Zakharov and Filonenko [1967a,b] (section A.7). They are important for systems in which the dissipation sink can absorb arbitrary rates of energy. Remarkably they are realized as a nontrivial power law spectrum *n*^KZ^ ∝ *k*^*v*^, with *v* < 0, *v* ≠ −1. The slope *v* of the spectrum is a value that reflects the dimensionality of the system, as well as its linear nonlinear properties (homogeneity degrees of the interaction coefficient and dispersion relation).

## 3.4. A demonstration of the weak turbulence ansatz: dynamics and kinetics of a single triad

Some features of the high-speed bispectra are consistent with stationary solutions of the dynamical equations 8. We illustrate this using a simplified, universal toy model derived from the ansatz dynamical equation by considering a single triad of modes *k* = *κ*_1_, *k*_1_ = *κ*_2_, *k*_2_ = *κ*_3_, with *κ*_1_ + *κ*_2_ = *κ*_3_ (see equation 10). If interactions with all other modes are ignored (equivalent to a single blue dot in figure 7), equations 8 to reduce to three equations, called the “three-wave system” (section A.8; Craick 1985, Rabinovich and Trubetskov 1989). Written in amplitude/phase these are

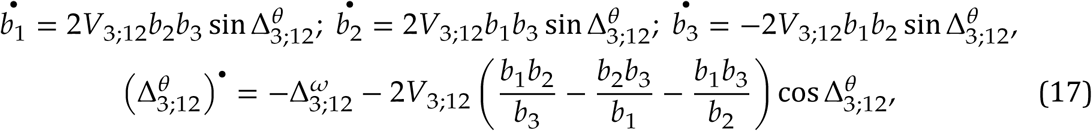

where *b*_*j*_ and *θ*_*j*_ are amplitudes of modes *κ*_*j*_, *j* = 1,2,3, and 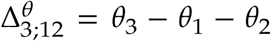. Averaged over realizations, the quantity 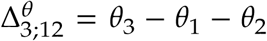 may be identified with the biphase (see section 3.1.2) If the triad is resonant, the kinetic version of the three wave equation is [e.g., Rabinovich and Trubetskov, 1989]

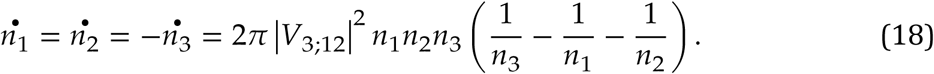

A large body of literature is available that investigates the relevance and dynamics of single-triad interactions in many physical situations, including plasma physics [Coppi et al., 1969, Weiland and Wilhelmsson, 1977, Craick, 1985], nonlinear optics [Ablowitz and Segur, 1981, Boyd, 2003], internal oceanic waves [Phillips, 1977, Craick, 1985], and other fields.

One may readily check that a stationary solution of equations 17 is given by the conditions: 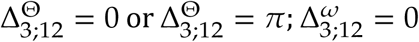 (the triad is resonant, equations 16), and that

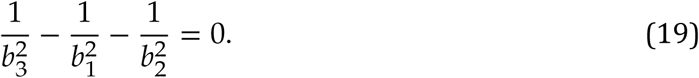

The last equation has, for example, the trivial solution 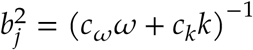 (RJ spectrum, thermodynamic equilibrium). In other words, stationary state solutions of the ansatz equations exhibit naturally 8 a biphase of 0 or *π*.

Applied to the triad formed by theta and its first harmonic (*θ,θ*,2*θ*), i.e., *κ*_1_ = *k*_*θ*_, *K*_2_ = *k*_*θ*_, and *K*_3_ = *k*_2*θ*_ (red dot in figure 7) this result is consistent with observations that show theta and in first harmonic are in phase (see figure 4b, upper-right panel). The effect of this type of coupling is to sharpen the crests and flatten the troughs of the time series (positive skewness, see figure 8, left panel).

**FIGURE 8.**
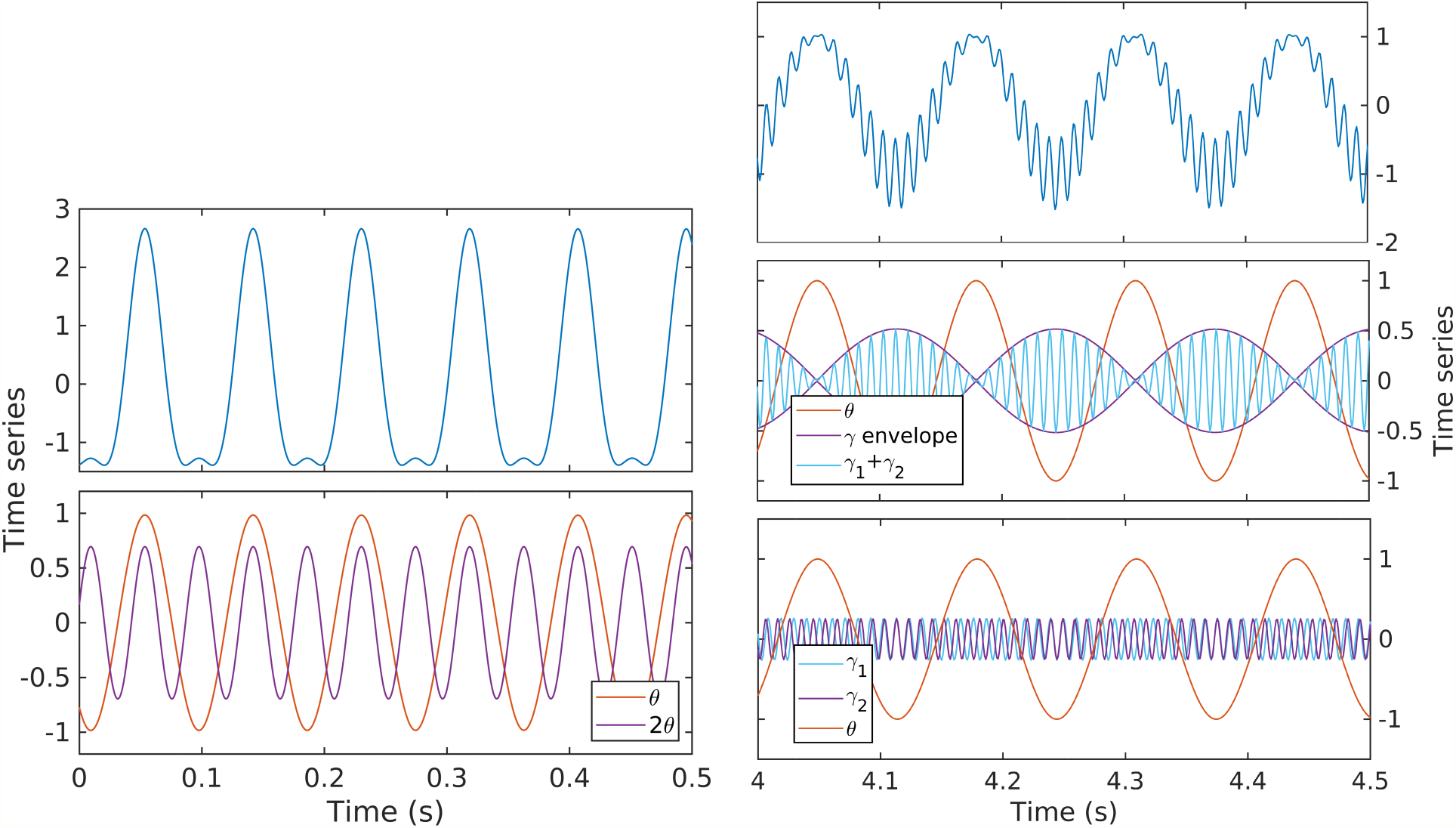
Examples of numerical integration of the three wave equation 54 for stationary states, under the assumption of resonance (equations 16), with *b*_*j*_ satisfying equation 19. Upper panels: total signal. Time series units are arbitrary.

**FIGURE 9.**
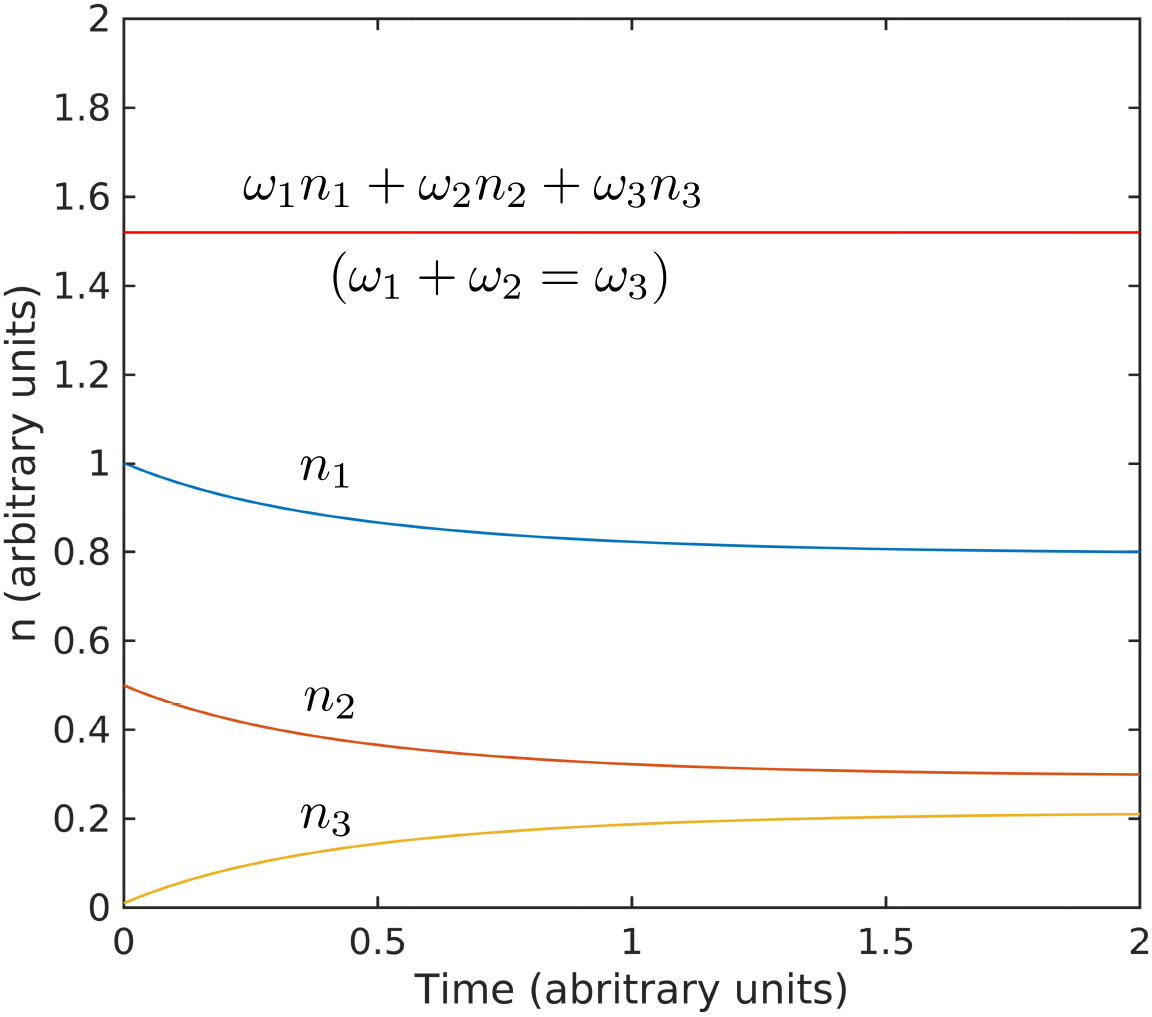
Box initially at rest on sled sliding across ice.

Applied to a theta-gamma triad (*γ, θ, γ + θ*), *κ*_1_ = *k*_*θ*_, *κ*_2_ = *k*_*γ*_, and *K*_3_ = *k*_*θ*_ + *k*_γ_ (yellow dot in figure 7), this result is consistent with the biphase value of *π*in figure 4b, upper-right panel. It is easy to check that effect of this type of coupling is that gamma envelope is maximal in theta troughs. Indeed, let 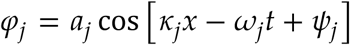, where *k*_*j*_ and *w*_*j*_ satisfy the resonance conditions 16. Elementary trigonometric manipulations yield the gamma envelope ∝ 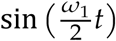. In other words, the gamma envelope is in quadrature with theta and its period is twice that of theta (figure 8, right panel). The kinetic evolution of a triad is illustrated in figure 9. The interactions described by the three-wave model result in time-reversible cyclic transfers of energy (the total energy is conserved if the system 54 is at resonance). In contrast, the long-time, *average* behavior (equation 18) shows a slow irreversible trend toward the stationary solution. If a full set of triads is taken into account, because all triads interact, a weak energy flow (turbulent cascade) develops that has the effect of driving the system of modes toward stationary distribution of energy over scales (RJ spectrum). If a sink is introduced in the small scales (e.g., high frequencies), the flow will naturally be directed toward the leaking mode, where it is taken out of the system. In this case a stationary state can be maintained only if energy is pumped into the system at the leakage rate, and a stationary state is realized if the energy is injected by the source at the rate it is lost to the sink. The cross-scale flow of energy is in this case constant at all scales. This type of stationarity corresponds to the KZ spectra.

Evolution toward stationarity of the solution of the kinetic equation 18. The initial conditions used *n*_1_(0) = 1, *n*_2_(0) = 0.5, and *n*_3_(0) = 0.01, and interaction coefficient are arbitrary and exaggerated to highlight the behavior. The Hamiltonian *w*_1_*n*_1_ + *w*_2_*n*_2_ + *w*_3_*n*_3_ is conserved for *w*_1_ + *w*_2_ = *w*_2_ (values for *ω* are also arbitrary).

## 4. DISCUSSION

While a wealth of knowledge has accumulated in recent years about brain activity at microscopic and macroscopic scales, the need for a consistent theory of the dynamics of intermediate-scale (mesoscopic) processes has received comparatively little attention, in part because, in some parts of the nervous system, they are the expression of activity of complex microscopic structures (e.g., for example central pattern generators can maintain specific, yet re-configurable rhythms; e.g., Rabinovich et al., 2012, Gutierrez et al., 2013, Marder et al., 2016). Although some microcircuits, with definite structure, may be able to impose an oscillation on the mesoscale, the cortex, with an amorphous structure, is rhythmically organized in a different manner, with a different source and might play a different role.

A number of previous studies [Lashley 1942, Hebb, 1958, Freeman, 2000b, 2007, Freeman and Vitiello, 2010] hypothesize that mesoscopic processes in the cortex represent the essential cognition step of abstraction and generalization, and therefore provide an essential mechanism for integration of brain activity at all scales. This observation is particularly intriguing, because the amorphous structure of the cortex at the anatomical mesoscale (e.g., any highly recurrent region in which activity is projected back into the same region or dense inter-connectivity mediated by interneurons; Lorente de No, 1938, Freund and G., 1996, Buzsaki et al., 2004, Mante et al., 2013) suggests that the material support of activity in the temporal mesoscale (e.g., gamma frequency band) is collective neural activity, Freemanss [2000b] “mass action”. These ideas suggest that the physics of mesoscopic collective action in the cortex is intimately related to cognition; that *the physics of collective action is, in fact, the physics of cognition*.

The focus of this study is the dynamics of mesoscopic collective action and its role in the general energy balance in the brain. Despite the potentially paramount importance of the topic, studies dedicated to its physics are few (but of outstanding quality; e.g., Wilson and Cowan, 1972,1973, Cowan et al., 2016, Wright and Liley, 1995, Troy, 2008). Here, we attempt to lay the foundation of a systematic theory of collective action. A few recent studies show that collective action in the hippocampus takes the form of propagating waves [Lubenov and Siapas, 2009, Patel et al., 2012, 2013, Zhang Honghui et al., 2018, Muller et al., 2018], but in general, information about its spatio-temporal organizatione is scarce.

The discovery of the strong correlation between rat speed during active exploration and hippocampal activity provides a parametrization of the evolution of hippocampal activity with behavior. Our observations of scale distribution of LFP power (spectra) and the leading order estimators of cross-phase coupling of LFP oscillations (bispectra) in the lacunosum molecu-lare layer show a strong and remarkably ordered evolution (representative of both CA1 and dentate gyrus layers). The LFP power and phase coupling in the theta and gamma frequency bands increases consistently with speed. The lowest levels of LFP power are associated with a featureless power law distribution. The high-power spectra exhibit distinctive spectral peaks at theta frequency and harmonics, and significantly increased levels of gamma activity. In the transition, at intermediate states, the spectrum tilts to a smaller-slope shape that extends progressively into the gamma range generating the appearance of a spectral front. The existence of a non-zero energy, lowest-level spectrum in the absence of coherent collective action is strongly suggestive of a self-organized critical background state [Buzsaki, 2006]. As collective action energy increases with exploration, the decreasing spectral slope, the appearance of a spectral front, and the development of a broad gamma peak, are strongly suggestive of a transfer of energy from the low frequencies (large scales) toward high frequencies (microscopic scales; [Buzsaki and Draguhn, 2004]). The similarity of collective action with the general turbulence theory is striking. This motivated us to propose weak turbulence as an ansatz for collective action dynamics.

In summary, we propose that the mesoscopic hippocampus may be described as an amorphous (homogeneous, isotropic), active network containing a macroscopic number of randomly and densely connected neural units (neurons). Collective action represents a perturbation of a background state of the active network, that may be represented as a self-organized critical state. Because collective action is macroscopic with respect to neural units, we postulate therefore that it is fundamentally stochastic (the precise state of a single unit does not matter), weakly nonlinear, and weakly dissipative. These form a minimal set of features for the development of a turbulence theory of collective action. We summarize the principles of the turbulence ansatz and demonstrate its applicability to observations by showing that the reduced (universal) three-wave interaction toy model provides results consistent with observations.

Our proposed description provides a unified view of the physics of active networks that reconciles the theories of self organized criticality and turbulence in the hippocampus and perhaps other regions of the cortex.

The turbulence ansatz has important consequences. As an immediate gain, it provides a new theoretical platform for developing data collection and analysis procedures, and a new framework for the interpretation of observations. Because the turbulence theory is well established, the interpretation of the results is straightforward, and their statistics is well understood. The central idea of the turbulence theory is the energy cascade through the inertial range of scales. Collective action physics appears to be consistent with the energy cascade concept [Richardson, 1922, Kolmogorov, 1941, Zakharov et al., 1992, Newell et al., 2001, Nazarenko, 2011]. In particular, the spectral tilting (slope change) observed in the evolution from the background state to high-speed states suggests a transition from self-organized criticality to some type of stationary turbulent state of the Kolmogorov-Zakharov kind [Zakharov and Filonenko, 1967a,b, Zakharov et al., 1992, Newell et al., 2001]. Further research is needed to understand the structure of this evolution. A systematic exploration of hippocampal dynamics starting from first-principles governing equations is ongoing.

Remarkably, the amorphous, recurrent mesoscale structure appears to be a general theme that is used across species and brain regions [Lorente de No, 1938, Parent and Hazrati, 1995, Marder and Bucher, 2001, Garamszegi and Eens, 2004, Apps and Garwicz, 2005, Mante et al., 2013], which suggests a “universal computational principle” with a comprehensive reconfiguration potential, especially under *a priori unknown* conditions[Sussillo and Abbott, 2009]. The nature of this computation process is not well understood. As often argued [e.g., Freeman, 2000a, Edelman and Gally 2001, Frisch, 2014], cognition processes cannot resemble a human-engineered system, built based on principles such as maximum simplicity, well-defined internal interactions, explicit assignment of function, no irrelevancy, and no adventitious compensation for error. Rather, they are expected to resemble biological systems: no design, no *a priori* function, and for which irrelevance has no meaning [Edelman and Gally 2001]. As Frisch [2014] states it, “biological systems have an intrinsic ability to maintain functions in the course of structural changes”, such that “specific functions can obviously be constituted on the basis of structurally different elements, a biological property that is referred to under the term degeneracy [Edelman and Gally 2001]”.

This begs the question, if collective action is fundamental for cognition, what is its role? We conclude this study by suggesting a possible answer. An intriguing paradigm of the computational function of mesoscale turbulence is offered by Liquid State Machines (LSM) models [Maass et al., 2002, Jager, 2002]. LSMs are online neural network models that process a time-windowed signal in real time. Their basic function is to perform a nonlinear transformation of the input, e.g., expand it into a wave field, and hold this information for a short duration, while output neurons extract local information from the field. Learning is achieved at the readout stage. Fading memory and input separability imply that LSMs are universal function approximators, and can serve as effective online classification pre-processor for the readout neurons. If the brain is a prediction machine, scrambling to assign meaning to streams of data^11^ in real time, processing of fragmentary information (short-time windows) is crucial. A LSM may perform fast, online, short-term memory pre-processing; learning is performed by long-term memory readouts, that can record optimal responses. It is conceivable that the hippocampus uses collective action in a way similar to a LSM, perhaps as a fast online classification machine, or as a dynamical system simulator. It seems plausible to imagine the cortex as a network of LSMs. In the least, the concept seems to agree with most natural systems.

## Conflict of Interest Statement

The authors declare that the research was conducted in the absence of any commercial or financial relationships that could be construed as a potential conflict of interest.

## Author Contributions

AS and AM collaborated closely in developing the turbulence ansatz for brain activity. AS developed part of the numerical methods and the Matlab® tools used fort the data analysis. YZ and YQ developed part of the numerical methods, and performed the numerical analysis. JPK managed data collection. AM provided the impetus for the research; designed, organized, and supervised the data collection; and provided the guidance and advice regarding all things neuroscience. AS wrote the manuscript with input from all authors.

## Funding

This work was supported by the McKnight Brain Research Foundation, and NIH grants- Grant Sponsor: National Institute on Aging; Grant number: AG055544 and Grant Sponsor: National Institute on Mental Health; Grant Number: MH109548.

## Acknowledgments

Special thanks to S. Burke for multiple insightful discussions, advice on the manuscript redaction and for patiently reviewing manuscript drafts; and to S.D. Lovett for technical support. AS is grateful to Prof. Victor Shrira for helpful discussions and suggestions.

## Data Availability Statement

The data sets for this manuscript are not publicly available because these data are part of ongoing student dissertation research.

## APPENDIX A. WAVE TURBULENCE ANSATZ: A SKETCH OF THE ALGEBRA

This is a summary of wave turbulence principles, abbreviated and simplified to retain only the main ideas. Turbulence is a rich theory, with deep implications for the physics of large systems, covering a wide range of topics that include hydrodynamics, plasma physics, nonlinear optics, aggregation-fragmentation processes, ocean waves, and many others. The goal of this succinct account is to provide a possible blueprint for future investigations in to the dynamics of mesoscale neural collective action.

A.1. **Governing equations**. Following the standard thermodynamics formalism [e.g., Callen, 1960], assume that the physical system is a one-dimensional spatial network whose mesoscopic state is completely described by a function *ϕ*(*x,t*) that represents the deviation from an appropriately chosen equilibrium state [e.g., Wright and Liley 1995]. Quite generally, we will assume that *φ* satisfies a weakly-nonlinear, non-dissipative evolution equation that tends is linear as *ϕ* → 0, i.e.,

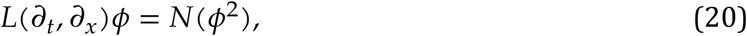

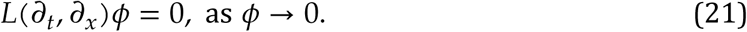

where *L* and *N* are constant-coefficient linear and nonlinear operators in *ϕ* and its derivatives. Because the nonlinearity is weak, if *ϕ* is not too large, we can neglect nonlinear terms *ϕ*^m^ with *m* > 2. Ignoring boundary conditions, equation 20 is solved using the Fourier transform

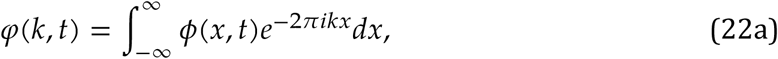

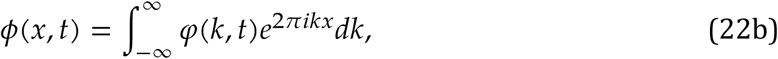

where *k* is the wavenumber, and the functions *e*^2πikx^ are orthogonal in the sense that

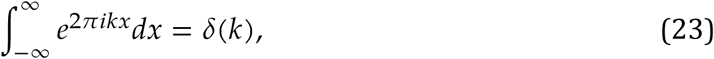

with *δ* the Dirac delta function, satisfying the sifting property 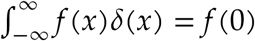. Infinitesi-mally close to the equilibrium state, substituting *φ*(*k,t*) = *A*(*k*)*e*^*-2πift*^ into equation 21 yields the dispersion relation between the frequency *f* and wavenumber *k* [Whitham, 1974]

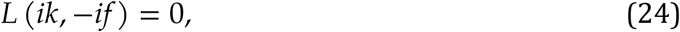

which may be solved to obtain *f* = *f*(*k*). The roots of the dispersion relation 24 are called modes. Neglecting dissipation implies that both *f* and *k* are real (with at most negligible imaginary parts). The only assumption we make about the dispersion relation is that the function *f*(*k*) (or *k*(*f*)) is monotonically increasing for positive wavenumbers and frequencies (higher frequencies correspond to higher wavenumbers and smaller scales, i.e., shorter waves).

A.2. **Hamiltonian formalism**. A plausible way to introduce a Hamiltonian description is as follows. Assume that the function *ϕ* may be expressed as as function of *r* extensive state variables, *ϕ* = *ϕ*(*q*), where *q*(*x,t*) = (*q*_1_,*q*_2_,…*q*_*r*_)(*x, t*). The function *ϕ* might be related to the local electrical field potential, and state variables might be physical space densities that describe mesoscale activity, such as number of neuronal firing pulses per unit network length, number of excitatory pulses received per unit network length, and so on. If *ϕ* completely characterizes the thermodynamics of the system, it determines all relevant intensive variables *p*(*x, t*) = (*p*_1_,*p*_2_, …,*p*_*r*_) (*x, t*), through the standard thermodynamic relations

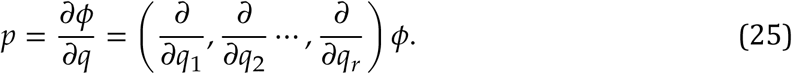

Let

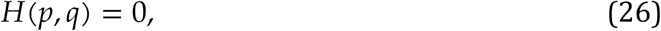

be an equation of state, where *H* is a function of the extensive/intensive thermodynamic parameters *q* and *p*. Equation 26 is a partial differential equation for *ϕ* (by substitution of *p* from equation 25). One can readily verify by substitution that the solution to equation 26 is given by the equations

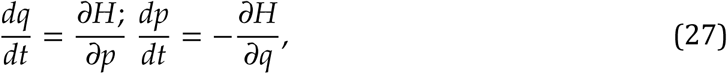

where *t* parameterizes the evolution of the system. Equations 27 maybe interpreted as Hamilton’s canonical equations, describing the evolution of the system in the space defined by the generalized coordinates *q* and momenta *p*, and subject to the constraint *H* = 0, where *H* is recognized as the Hamiltonian of the system [e.g., Peterson, 1979, Rajeev, 2008, Bal-diotti etal., 2016]. A change of variables (*p,q*) that preserves the form of the canonical equations 27 is called a canonical transformation. The symmetry of the Hamiltonian description may be used to further simplify the dynamical equations by performing two canonical transformations: the so-called Bogoliubov transformation [e.g., Zakharov et al., 1992] (*q*(*x,t*), *p*(*x,t*)) → *a* (*k,t*)

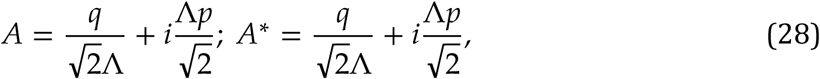

where the constant Λ is used to bring the variables *q* and *p* to the same physical units, followed by a Fourier transformation (the transform 22a-22b is canonical and unitary) *A*(*x, t*) → *a*(*k,t*), where

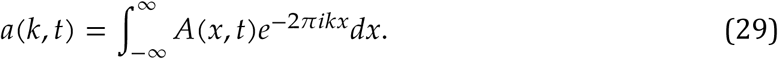

In the new variables, the general form of the Hamiltonian corresponding to equation 20, without creation-anihilation terms, [e.g., Zakharov et al., 1992], and retaining only the leading order nonlinearity, is

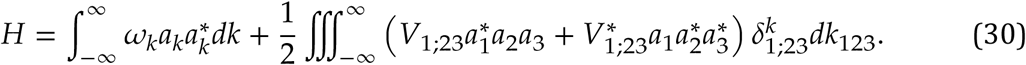

To simplify the notation we used the following shorthand notations: *a*_*j*_ = *a* (*k*_*j*_, t), *w*_*j*_ = 2*πf*(*k*_*j*_); and *a*_*k*_ = *a*(*k,t*),*w*_*k*_ = *w*(*k,t*) for modes *k* and *k*_*j*_,;*dk*_23_ = *dk*_2_*dk*_3_, 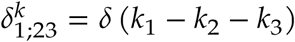. The nonlinear term is a convolution integral of all the nonlinear terms. The interaction coefficients *V*_1;23_ = *V*(*k*_l_,*k*_2_,*k*_3_) depend on the wavenumber, and are symmetric in the indices 2 and 3. If the Hamiltonian is identified with the energy of the system (conserved), the quantity |*a*|^2^ has dimensions of action (energy × time).

The Hamiltonian form 30 is universal; the details of the physics of the system are contained in the dispersion relation *w* = *w*(*k*) and the structure of the interaction coefficient V_1;2_3. The canonical equations become

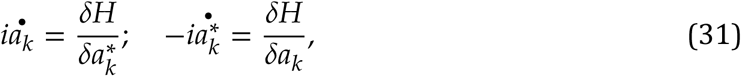

where we used the bullet notation for time derivative 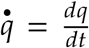 and introduced the standard notation *δ* for the variational derivative. Because the two equations are obtained from each other through complex conjugation, the system 27 is now reduced to equation a single equation (the second equation is simply its complex conjugate). Substituting the Hamiltonian 30 into equations 31 obtains equation 8. Equation 8, usually referred to as the dynamical equation, is the basis of our ansatz, and the main object of this discussion. Under the assumptions made so far, like the Hamiltonian form 30, equation 8 is universal, with the physics of the system contained in the coefficients.

A.3. **The BBGKY hierarchy and the kinetic equation**. The goal of averaging of dynamical equation 8 is to derive evolution equations for moments of the probability distribution of *φ*, or alternatively, its cumulants. In the Fourier space, this is equivalent to deriving the evolution equations for quantities known as “correlators”, such as 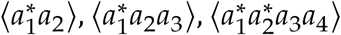, and so on, where the angular brackets denote the ensemble average. While the derivation is of the equation is straightforward, the resulting system is comprised of an infinite sequence of equations that, at each order, involve correlators of higher order, e.g.,

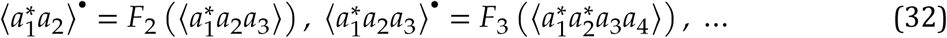

and so on, where *F_2_* and *F_3_* are some functions. System 32, known as the BBGKY hierarchy (Bogolyubov-Born-Green-Kirkwood-Yvon; e.g., Montgomery and Tidman, 1964, Alexeev, 2004), is not closed and cannot be solved, unless some means of truncating it (closure) are found. The closure problem is familiar to statistical mechanics. We provide here a sketch of the calculations, following procedures detailed in Newell, 1999, Newell et al., 2001, Zakharov et al., 1992, Zakharov, 1999, Nazarenko, 2011 and others.

Assuming spatial homogeneity implies that The quantity *n (k)* represents the action density, but is also referred to as “occupancy number” or “number of particles”, by analogy with quantum mechanics. We will also call *n* and *ℱ* by their generic stochastic-process names of “spectrum” and “bispectrum”, respectively.

A.3.1. *Double correlator (spectrum)*. To derive an equation for the double correlator multiply equation by 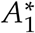 and subtract from it its complex conjugate and average using the spatial homogeneity assumptions (equations 11- 12) obtains the lowest order equation of the BBGKY hierarchy,

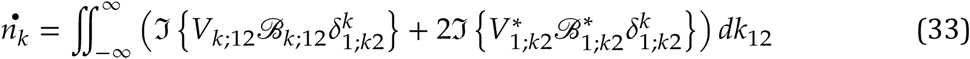

describing the evolution of the spectrum *n* as a function of the bispectrum.

A.3.2. *Triple correlator [bispectrum)*. An equation for the evolution of the bispectrum *ℱ* can also be derived from the dynamical equation 8. Differentiating to time the triple products 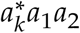 and 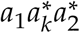 and averaging yields, for example for the first product in the equation for the double correlator

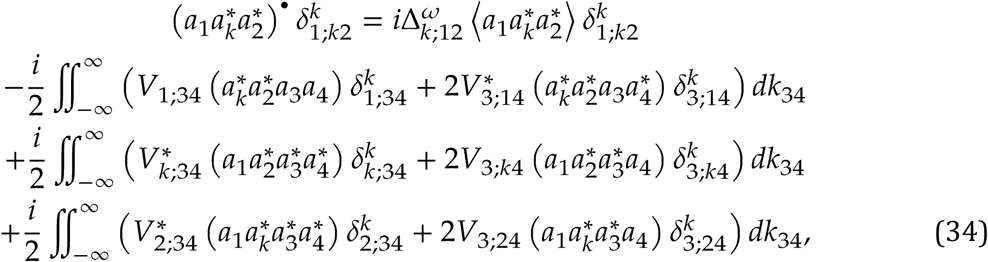
showing that the evolution of triple correlators is driven by quadruple correlators 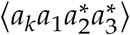. The same procedure can be applied to derive evolution equations for higher-order correlators. At each step next order correlators are involved, thus the resulting system of equations is not “closed” and cannot be solved. Assumptions that lead to the closure of the system have to be made.

A.4. **Quasi-Gaussian closure**. If amplitudes are small and stay small through the evolution process, say *a* = *O*(*∊*), where *∊* ≪ 1, then correlators are (and stay) well ordered, i.e., *a*_1_*a*_2_) = *O*(*∊*^2^), 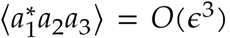,and so on. This property of the equation is commonly referred to as weak nonlinearity. Then fourth-order correlators 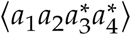 should have a generic quasi-Gaussian structure (i.e., dominated by variance, e.g., Newell et al. 2001, Nazarenko 2011),

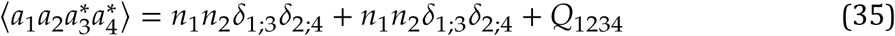

where Q_1234_ is a irreducible residual of higher order (∊^−5^). Substituting equation 35 into the average of the triple product in equation 34 and neglecting the irreducible terms yields the equation for the evolution of the bispectrum

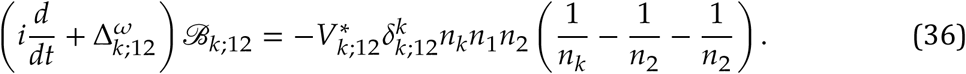

Equations 33 and 36 form the system of coupled equations 13. The BBGKY system is closed, since the evolution of the bispectrum depends only on the spectrum. Equations 13 describe the stochastic evolution of the system, on time scales of order *O* (∊^−4^)

We should stress that this closure is valid only for as long as the correlators are well ordered. If singularities appear as a result of the evolution, the closure breaks down. Simple scaling consideration [Newell and Rumpf, 2010] show that the closure is scale dependent and necessarily breaks a small scales.

A.5. **The kinetic equation**. With some standard simplifications [e.g., Zakharov et al., 1992, Zakharov, 1999, Anenkov and Shrira, 2018], the system 33-36 may be simplified further. Assuming that the spectrum varies with time much slower than the linear phase (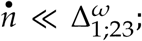 averaging the modulus squared 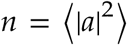 eliminates the fast oscillatory time-dependence), equation 13b can be integrated approximately for 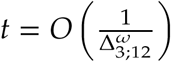 to obtain

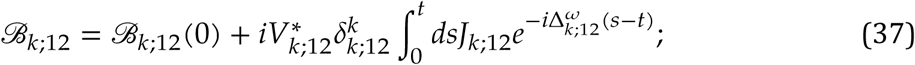

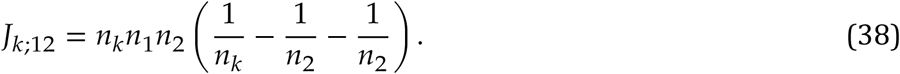

If the initial bispectrum is zero (*ℱ*_*k;12*_ (0) = 0) and factoring out of the integral the expression *J*_*k*;12_ containing the spectra obtains

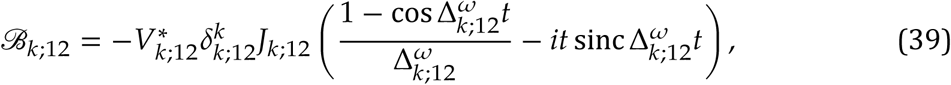

where 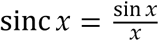. The longtime limit *t* → ∞ of equation,

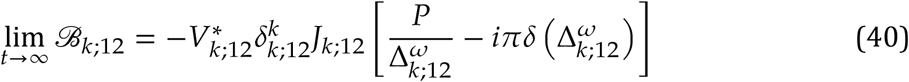

where 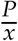 is Sokhozki’s generalized function [Vladimirov, 2002], satisfying the relation

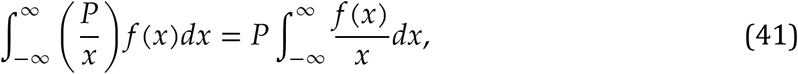

with *P* denoting the principal value of the integral. This solution is meaningful only if 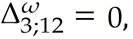 a condition that comes in addition to the selection criterion 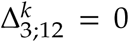 for the triad. The system of equations

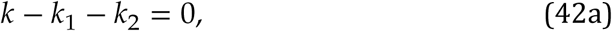

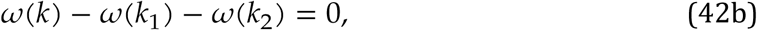

is called “resonance conditions”.

Substituting into the equation for the evolution 13a of the spectrum, which requires only the imaginary part of the bispectrum obtains, after symmetrization, equation 14, the so-called kinetic equation.

A.6. **Conservation laws for the kinetic equation**. The dynamical equation 8 has one integral of motion, the Hamiltonian 30. The kinetic equation 14 has its own set of integrals. Let Q be the *physical-space* density of an extensive quantity,

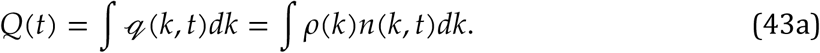

where *ρ(k)* does not depend on time. Examples of such quantities are the energy *E* with densities *e* = *ω(k)n*, and momentum *M* with densities *m* = *kn.* Using the kinetic equation, the time derivative of *Q* is

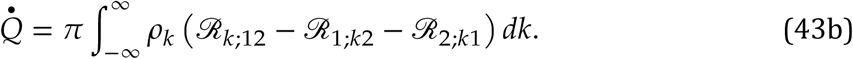

The quantity *Q* is conserved if 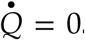 In gWeneral, the conservation equation 44 may also be recast as a continuity (transport) equation

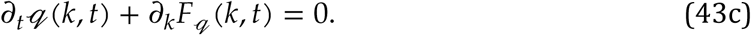

where 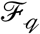 is the spectral flux of 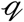. Comparing equations 44 and 43c, one can write

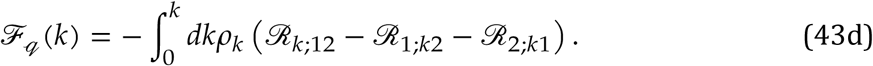

If the density *q(k,t)* is stationary then 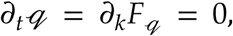 *Q* is conserved, and the spectral *q*-flux is constant.

Such conserved quantities exist. For example, if |*V_k;12_*|^2^ is invariant to permutations of indices, relabeling in equation 43b 1 ↔ *k* in 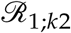 and 2 ↔ *k* in ↔ 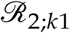 transforms it into

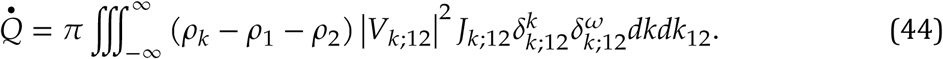

The product 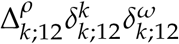 cancels if *ρ* = *ω(k)* or *ρ* = *k.* It is obvious that the energy *E* and momentum *M*, defined as

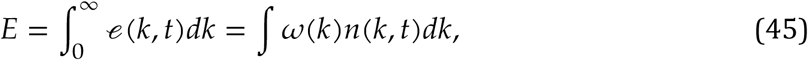

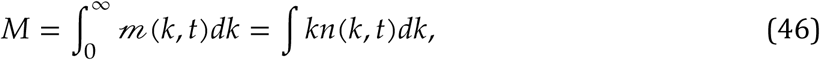

are conserved by the kinetic equation.

A.7. **Stationary solutions of the kinetic equation**.

A.7.1. *Thermodynamic equilibrium, Rayleigh-Jeans (RJ) spectra: A* family of stationary spectra for equation 14 is immediately found by inspection. If we set

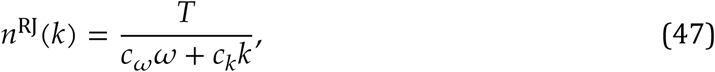
where *T,c*_*ω*_ and *c_k_* are constants, the integrand 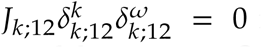 for *all triads* - Moreover, this spectrum corresponds to the equipartition of the quantity *Q* with density *q*(*k*)=(*c*_*ω*_ω+*c*_*k*_*k*)*n*_*k*_ for example, if *c*_*k*_ = 0, this represents the equipartition of energy. This type of stationary state corresponds to a “detailed balance”, where nonlinear interaction cancels for each triad, and 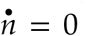 regardless of the quantity *q* carried by the number of particles *n.* The spectral *q*-flux cancels, in equation 43c *F*_*q*_ = 0.

Thermodynamic equilibrium states cannotbe realized at all scales because equipartition means that the *physical-space* density *Q* = ∫ *Tdk* is infinite (a phenomenon known as the ultraviolet catastrophe). However, it might occur if the spectral fluxes “stagnate” near a critical wavenumber *k_0_*, preventing fluxes to infinite wavenumbers. The system will tend to “ther-malize” i.e., approach a zero-flux, thermodynamic equilibrium state (a discussion of this “bottleneck” scenario is given in Nazarenko, 2011).

A.7.2. *Kolmogorov-Zakharov spectra.* In non-isolated systems that have sources and sinks of *Q* well separated in the spectral domain, one would expect non-zero fluxes *F*_*q*_ from the sources to sinks, similar to the hydrodynamic energy cascade described by Richardson and Kolmogorov [Richardson, 1922, Kolmogorov, 1941, Frisch, 1995]. Nonzero stationary solutions of the kinetic equations were found by Zakharov and Filonenko [1967a,b] and are known as the Kolmogorov-Zakharov (KZ) spectra.

At stationarity, the integral on the left-hand side of equation 14 cancels. Under quite general conditions (e.g., if only one physical process is involved), the dispersion relation and the interaction coefficients are homogeneous of degree *α* and *β*, respectively:

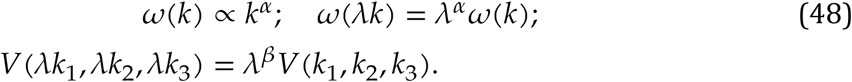

Because we are in a unidimensional system, we are not concerned is isotropy. Look for a solution in the power law form

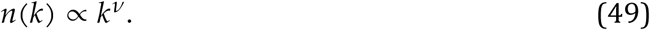

in equation 14, change the integration variables to (Zakharov transformation)

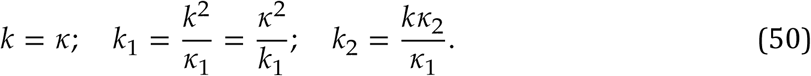

transforms the second to

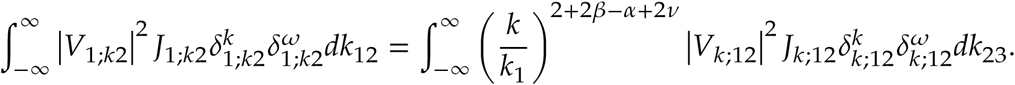

Applying a similar transformation to the third term and denoting *x* = −(2+*β* − *α*+ 2*v*),the kinetic equation becomes

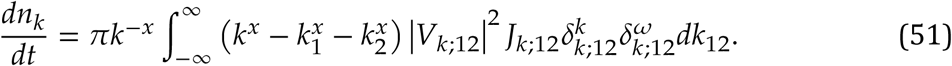

The spectrum *n*(*k*) is stationary if (compare with equation 44) either 1) *x* = *ω,v* = − 1 − *β*, corresponding to the conservation of linear energy, or if 2) *x* = 1, 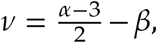 corresponding to conservation of momentum. Neither is an RJ spectra, since *v* ≠ *α*. and *v* ≠ 1, therefore neither cancels the integrand *ℛ*_*k*;12_. Note that because the Zakharov transformation 50 is singular in *k*_1_ = 0 (non-identity transformation) and the integrals in the kinetic equation 14 are not bounded, the validity of the KZ stationary solution is not guaranteed unless the convergence of the integrals is verified.

A simple scaling argument may be used to show that the spectral energy flux is constant in the the case *x* = *α* The continuity equation 43c for energy (equation 45) in the spectral domain is

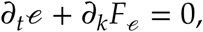

where the energy flux is (see equation 43d)

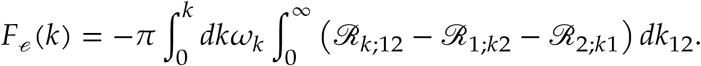

Scaling all wavenumbers by *k*, i.e., changing variables (*k,k*_1_,*k*_2_) → *k*(*κ,κ*_1_,*κ*_2_) brings the spectral energy flux to the scaled form

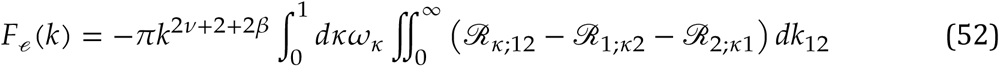

Setting the scaling factor to be independent of *k*; means setting 2*v* + 2 + 2*β* = 0, which obtains*v* = − 1 − *β*, i.e., the KZ spectral slope obtained above. A similar argument shows that thesecond KZ spectrum 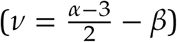 corresponds to constant spectral fluxes of momentum.

A.8. **The three-wave equation**. The three wave equation is a universal model [Weiland and Wilhelmsson, 1977, Craick, 1985, Zakharov et al., 1992] deriving from the dynamical equation 8 by restricting the interaction to a single triad of modes (*k*_1_, *k*_2_, *k*_3_) satisfying the selection criterion *k*_3_ = *k*_1_ + *k*_2_

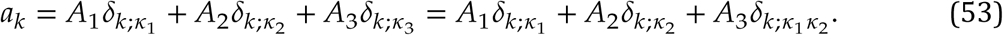

Substituting equation 53 into the dynamical equation 53 obtains equation

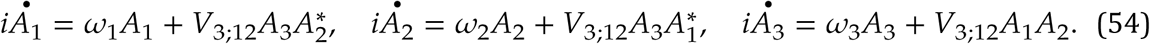

Substituting *A*_*j*_ = *b*_*j*_*e*^*iθ*_*j*_^, with *b_j_*, *θ_j_* ∈ ℝ and *b_j_* > 0, obtains the amplitude-phase representation

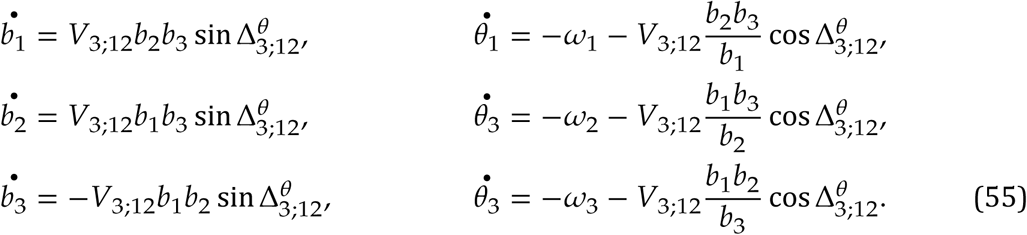

The system 55 may be reduced to 4 equations and has analytical solution given in terms of elliptic functions. Combining the last three equations yields a system of four equations with four unknowns: the amplitudes *b*_*j*_, *j* = 1,2,3, and the phase 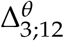

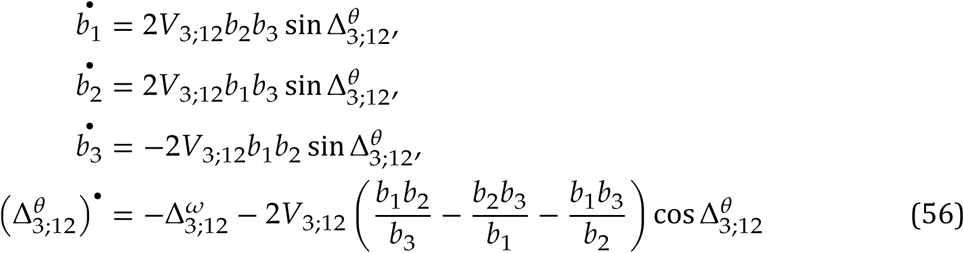

Following the procedure outlined in section A.3, and assuming the triad is resonant (see equations 42) one obtains the kinetic three-wave equations [e.g., Rabinovich and Trubetskov, 1989]

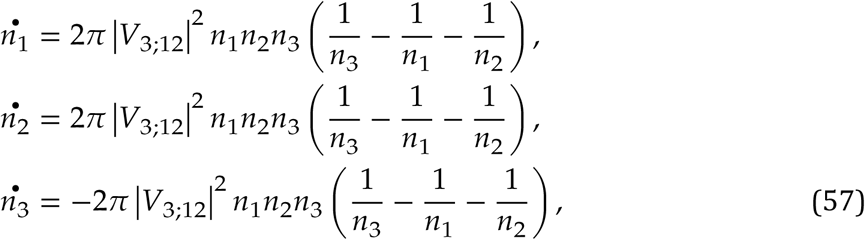

where the notation is the same as the one used for the full kinetic equation 14. Because the physical system described by equations 57 comprises only one triad, stationarity conditions degenerate to detailed balance. The RJ spectrum 47 is obviously a solution: direct substitution of expression 47 into equations 57 cancels the factor in parentheses. Energy and momentum are conserved regardless of whether are stationary or not, because the triad is resonant (equations 42)

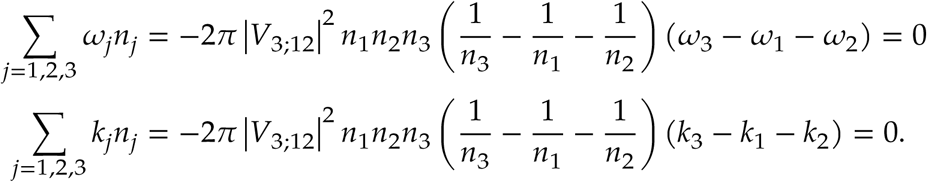

## Footnotes

1 Frequency bounds given here are just convention, and could be modified depending on the processes examined.

2 While action potentials are the fundamental process that carry the signal, like any natural process, it burns more energy than it uses. Stated differently, it is not a lossless system. Rather, synaptic transmission and action potentials comes at the price of ion exchange/energy loss (ATP required to maintain membrane charge).

3 The amorphous (homogeneous and non-isotropic character of the hippocampal CA1 is illustrated by the randomness and density of connections: ~390,000 neurons [Witter and Amaral, 2004], with approximately 30,000 putative excitatory synapses and ~1700 inhibitory synapses onto a single pyramidal cell [Megias et al., 2001], also connected to interneurons [Freund and G., 1996, Marshall et al., 2002] which in turn contact several hundred principal neurons [Sik et al., 1995].

4 The phrase “collective activity” is equivalent to Freemans [1975] “mass action” concept. We prefer “collective” over “mass”, because the word “mass” has a reserved meaning in physics.

5 Freeman and Vitiello [2010]: The problem was clearly stated over fifty years ago: “Generalization is one of the primitive basic functions of organized nervous tissue. Here is the dilemma. Nerve impulses are transmitted … from cell to cell through definite intercellular connections. Yet all behavior seems to be determined by masses of excitation. … What sort of nervous organization might be capable of responding to a pattern of excitation without limited specialized paths of conduction? The problem is almost universal in the activities of the nervous system [Lashley, 1942]”.

6 Discussing strategies for optimizing the use of large data sets O’Leary et al. [2015] note that“… there is still a big role for conceptual models that tell investigators what kinds of processes may underlie the data, or, more importantly, what potential mechanisms one should rule out”.

7 The reader might find the history of the debate surrounding the work of Boltzmann [1872, 2003] and the birth statistical mechanics quite instructive (e.g., Pathria and Beale, 2011).

8 Other scenarios are easy to imagine and have been observed. For example, if the dissipation capacity off the sink is smaller than the input rate at the source, energy will accumulate in the small scale range, possibly causing some system failure (e.g., a switch to different physical regime). This is a type of bottleneck scenario [e.g., L’vov et al., 2007, Meyers and Meneveau, 2008, Proment et al., 2009, Nazarenko, 2011].

9 Scale localization of dissipation processes is common in physics. In the case of water waves, for example, long swells (ocean waves people surf; wavelength ~100 m) lack a direct dissipation mechanism and can propagate for hundred of kilometers with negligible decay, while capillary waves (wavelength ~ cm) experience strong dissipation. Because energy cannot jump scales from swells directly to capillary waves, swells decay by cascading their energy into progressively smaller scales until the dissipation scale is reached.

10 Scale localization of dissipation processes is common in physics. In the case of water waves, for example, long swells (ocean waves people surf; wavelength ~100 m) lack a direct dissipation mechanism and can propagate for hundred of kilometers with negligible decay, while capillary waves (wavelength ~ cm) experience strong dissipation. Because energy cannot jump scales from swells directly to capillary waves, swells decay by cascading their energy into progressively smaller scales until the dissipation scale is reached.

11 In the “guess the phrase” game, the letters of a common phrase, initially blank, are filled out randomly; the goal is to identify the full sequence the fastest.

